# Dishevelled localization and function are differentially regulated by structurally distinct sterols

**DOI:** 10.1101/2024.05.14.593701

**Authors:** Sonali Sengupta, Jazmine D.W. Yaeger, Maycie M. Schultz, Kevin R. Francis

**Affiliations:** Cellular Therapies and Stem Cell Biology Group, Sanford Research, Sioux Falls, SD, 57104, USA; Department of Pediatrics, University of South Dakota Sanford School of Medicine, Sioux Falls, SD, 57105, USA

**Author notes:** **Corresponding author. Requests should be addressed to K.R.F. (**).

**Keywords:** Dishevelled, Wnt, cholesterol biosynthesis, sterols

## Abstract

The Dishevelled (DVL) family of proteins form supramolecular protein and lipid complexes at the cytoplasmic interface of the plasma membrane to regulate tissue patterning, proliferation, cell polarity, and oncogenic processes through DVL-dependent signaling, such as Wnt/β-catenin. While DVL binding to cholesterol is required for its membrane association, the specific structural requirements and cellular impacts of DVL-sterol association are unclear. We report that intracellular sterols which accumulate within normal and pathological conditions cause aberrant DVL activity. *In silico* and molecular analyses suggested orientation of the β- and α-sterol face within the DVL-PDZ domain regulates DVL-sterol binding. Intracellular accumulation of naturally occurring sterols impaired DVL2 plasma membrane association, inducing DVL2 nuclear localization via Foxk2. Changes to intracellular sterols also selectively impaired DVL2 protein-protein interactions This work identifies sterol specificity as a regulator of DVL signaling, suggests intracellular sterols cause distinct impacts on DVL activity, and supports a role for intracellular sterol homeostasis in cell signaling.

## INTRODUCTION

Dishevelled (DVL) proteins (DVL1/2/3) are best known for the integration and propagation of Wnt signaling in cells and tissues (Gao and Chen, 2010; Sharma et al., 2018). Early genetic studies in Drosophila demonstrated the DVL-dependent regulation of Wnt signaling in segment cell polarity and embryonic development (Fahmy and Fahmy, 1959; Wallingford and Habas, 2005). The vertebrate homologues of DVL have also been implicated in essential cellular processes such as proliferation, migration, development, differentiation, and polarity establishment (Klingensmith et al., 1996; Pizzuti et al., 1996). DVLs relay cellular signals through both the canonical and non-canonical Wnt signaling pathways (Paclíková et al., 2021) in addition to signaling pathways such as GSK3β (Hajka et al., 2021), RYK (Lu et al., 2004), polar cell polarity/calcium signaling (Clevers and Nusse, 2012), and mammalian target of rapamycin (mTor) (Cai et al., 2006; Caliskan et al., 2023; Mak et al., 2005). DVL signaling is critical to vertebral development (Gray et al., 2009; Na et al., 2007; Tadjuidje et al., 2011). While DVL paralogs are differentially expressed across embryonic stages (Gray et al., 2009), *DVL2* is the highest expressing isoform throughout embryogenesis (Klingensmith et al., 1996). Mutation of DVL isoforms in animal models cause defects in the development of the cardiovascular tract, neural crest, skeleton and cochlear systems (Etheridge et al., 2008; Hamblet et al., 2002) while autosomal dominant mutations in DVL isoforms lead to Robinow syndrome, characterized by broad malformations within the skeletal system, urogenital system, and neurological deficits (Schwartz et al., 2021; White et al., 2015; White et al., 2016). Lastly, changes in DVL isoform expression and localization have been reported in cancers including glioma (Li et al., 2014), colorectal (Metcalfe et al., 2010), and breast cancer (Metcalfe et al., 2010; Rasha et al., 2023).

The conserved domains of DVL proteins play a pivotal role in regulating DVL signaling through molecular binding properties. DVL proteins contain three conserved domains, an amino-terminal Dishevelled-Axin (DIX) domain, a carboxy-terminal Dishevelled, EGL-10 and pleckstrin (DEP) domain, and a central <u>P</u>ost-synaptic density protein (PSD95), <u>D</u>rosophila disc large tumor suppressor (Dlg1), and <u>Z</u>onula occludens-1 (ZO-1) (PDZ) protein domain. While the DIX-domain mediates DVL oligomerization, signalosome formation, and Axin binding to mediate β-catenin destruction complex formation (Liu et al., 2011; Schwarz-Romond et al., 2007), the C-terminal DEP domain is essential for targeting DVL proteins to the plasma membrane and for assembly of functional signalosomes (Gammons et al., 2016; Pan et al., 2004; Wong et al., 2000).

While the PDZ domain mediates the majority of DVL protein-protein interactions that are crucial for cellular processes (Wong et al., 2000), it requires lipid interactions to elicit signaling. The PDZ domain of DVL was previously shown to require interaction with cholesterol to support canonical Wnt signaling (Francis et al., 2016; Sheng et al., 2014). Cholesterol is critical to the formation of liquid-ordered (*L_0_*) domains (i.e. lipid rafts or lipid nanodomains) to support transient protein-protein or lipid-protein complexes linked to membrane-cytoskeleton communication, receptor-ligand interaction, and receptor clustering (Anderson et al., 2021; Azbazdar et al., 2019; Simons and Toomre, 2000). Removal of cholesterol via methyl-β-cyclodextrin (MβCD) also disrupts lipid rafts and perturbs Wnt signaling (Riitano et al., 2020). We previously demonstrated that within the cholesterol biosynthesis disorder Smith-Lemli-Opitz syndrome (SLOS), characterized biochemically by accumulation of the sterol intermediate 7-dehydrocholesterol (7DHC) concomitant with cholesterol loss, impaired Wnt/β-catenin signaling is associated with reduced binding affinity between DVL-PDZ and 7DHC (Francis et al., 2016). While these studies demonstrate a clear role for cholesterol-DVL interactions in mediating Wnt signaling, the specificity of DVL-lipid interactions and the resulting impacts on DVL activity require elucidation.

In this study, we demonstrate that structural differences between cholesterol and distinct sterols with biological and clinical significance inhibit DVL membrane association via PDZ domain interactions. We also report that sterol changes redirect the subcellular localization of DVL2 and promotes nuclear translocation of DVL2. To our knowledge, this is the first causal link established between sterol depletion, sterol biochemistry, and DVL protein activity. Our work suggests that cell-specific and disease-specific sterol expression could directly impact DVL protein function with broad significance to developmental biology, tissue homeostasis, and human disease.

## RESULTS

### Molecular modeling and *in silico* docking suggest sterol structure impacts DVL membrane association

While previous studies have identified cholesterol binding motifs for transmembrane receptors and other membrane-integrated proteins (Bukiya and Dopico, 2017; Fantini and Barrantes, 2013; Marlow et al., 2021), the ability of membrane-associated proteins such as DVLs to bind cholesterol versus other sterols is not established. To test the hypothesis that sterol content and sterol structure regulate the membrane binding affinity of DVL proteins, we performed *in silico* ligand-receptor binding analysis between the PDZ domain of DVL2 and either cholesterol or selected sterols of biological and clinical relevance (**Figure 1A**; **Table S1)**. The chemical structure of cholesterol has several elements which regulate cholesterol-protein binding: the hydrophobic tetracyclic ring system, the C5-C6 double bond that confers rigidity, the flexible isooctyl side chain, and the polar C3 hydroxyl group. The alternate sterols selected exhibit changes in the position and number of double bonds, imparting variable rigidity among other features (**Figure 1A**).

**Figure 1.**
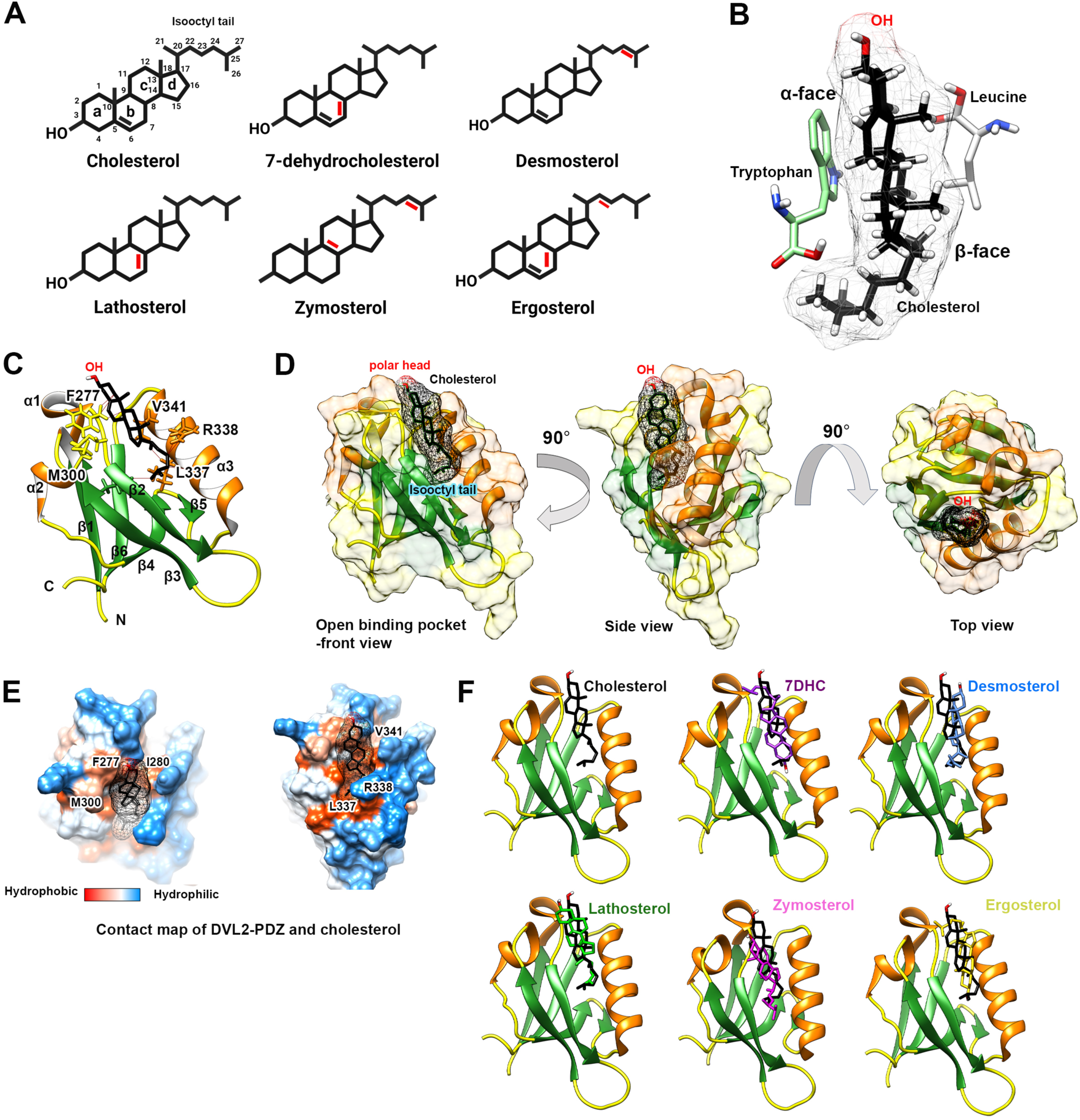
Homology modelling and receptor-ligand binding studies suggest DVL2 exhibits differing binding affinities for sterols. **A)** Chemical structures of sterols of interest (structural differences annotated in red). **B)** Favorable association of cholesterol (black with space-filling) with DVL2-PDZ domain amino acid residues (colors). While the smooth α-face of cholesterol favors interaction with an aromatic tryptophan, the rough β-face favors branched leucine interaction. **C)** Structure of the folded PDZ domain of DVL2 with bound free cholesterol (black). The secondary structures of the protein domain are annotated (helix, orange; strand, green; coil, yellow) as α1-3 and β1-5 from N-terminal to C-terminal. The predicted cholesterol:residue interactions (distance < 4 A°) are annotated (F277, I280, M300, L337, R338, V341). Protruding side chains of the corresponding residues are shown for visualization. Docked cholesterol is positioned longitudinally along an open binding pocket between α3 and β1. **D)** The space filling model of DVL2-PDZ with bound cholesterol showing the proposed binding pocket from front, side, and top view. **E**) Hydrophobicity calculation of the space-filling model of DVL2-PDZ with bound cholesterol. **F)** Differential binding of cholesterol (black) versus biologically important sterols (colors) to the DVL2-PDZ domain.

To perform differential binding analysis, the crystal structure of human DVL2-PDZ (PDB ID: 3CBZ) was used as the docking receptor **(Figure S1A).** 3-D conformers of sterols obtained from PubChem (https://pubchem.ncbi.nlm.nih.gov/compound/444679#section=2D-Structure) were used as ligands (**Figure S1B**). The 3-D model of the DVL2-PDZ domain presents three α-helices and six β-strands. When docked, cholesterol occupies a one-sided binding pocket formed between α, β2, and the floor of the cavity delineated by the β5/6 of DVL2-PDZ. While no hydrogen bonds were formed between the cholesterol and DVL2-PDZ, F277, I280, V341, R338, L337 and M300 surrounding the binding pocket were identified as possible interacting residues **(Figure 1B and 1C)**. Predicted contact residues were hydrophobic except R338, suggesting the binding pocket is likely hydrophobic and can accommodate the highly non-polar sterol (**Figure 1D and 1E**). Blind docking results suggested DVL2-PDZ exhibits different binding affinities for cholesterol versus selected sterols of interest (**Table S2)**, possibly due to altered sterol binding orientation in the hydrophobic pocket of the DVL2-PDZ domain (**Figure 1F**; **Figure S1**). There is poor consensus on the interaction motifs between cholesterol and soluble proteins. In the absence of hydrogen bonding, the transient binding is often a function of hydrophobic force between the sterol and protein (Reitz et al., 2008). In the DVL2-PDZ domain, the free cholesterol aligns itself along the longitudinally open pocket and the ligand is loosely aligned with hydrophobic patches on both sides of the tetracyclic rings with the isooctyl chain and the 3’ polar headgroup projecting outside (**Figure 1D and 1E**). This conformation across the α- and β-face supports previously observed characteristics of a cholesterol binding pocket in other proteins (Bukiya and Dopico, 2017). The highest scoring poses for other sterols reveal several degrees of rotation of the ligands along the DVL2 cleft with respect to cholesterol (**Figure 1E**). These predictive analyses suggest cellular sterol content may regulate DVL membrane localization and thus signaling.

### The membrane association of DVL proteins is regulated in a sterol-specific manner

While *in silico* analyses suggest differential binding of sterols to DVL2, these conformations assume an interaction between free cholesterol and DVL2, an unlikely physiological occurrence due to cholesterol being membrane bound and DVL2 being a soluble, transiently membrane-binding protein. To experimentally investigate the differential binding of sterols to DVL proteins, we performed dose-dependent plasma membrane (PM)-mimetic vesicle binding assays with purified recombinant *Hs*DVL2-PDZ domain peptide and other DVL variants (**Figure 2A; Figure S2A-C**). We purified the DVL2-PDZ domain as well as a 507 amino-acid long HsDVL2 displaying all major domains except the C-terminal tail (DVL2-FL; **Figure 2A**). We also purified DVL1 and DVL3, similarly truncated beyond the DEP domain (DVL1-FL, DVL3-FL, 474 and 496 amino acids, respectively) (**Figure 2A**). To model DVL association to the inner plasma membrane, unilamellar PM-mimetic vesicles (100 nm) were prepared to mimic the cholesterol and phosphoinositide concentrations in the cytoplasmic leaflet of the plasma membrane (Sheng et al., 2014). To compare DVL2-PDZ binding to cholesterol versus other sterols, cholesterol was replaced by equimolar concentration of alternative sterols within PM-mimetic vesicle preparations (**Figure 1A; Figure S2D-J**) followed by protein-PM-mimetic vesicle pulldowns with pellet fractions immunoblotted using anti-His antibody (**Figure 2B**). Relative binding efficiencies from western blots demonstrated that DVL2-PDZ binds cholesterol-PM vesicles most efficiently (>5-fold ergosterol, >2-fold 7DHC and zymosterol) (**Figure 2B, C**). Desmosterol and lathosterol exhibit DVL2-PDZ binding efficiencies similar to cholesterol (**Figure 2B, C**). To determine if these binding differences were DVL isoform specific, we also compared cholesterol and 7DHC binding to DVL1-FL and DVL3-FL (**Figure 2A**). For each DVL isoform, cholesterol-PM-vesicles exhibited higher binding efficiency than 7DHC **(Figure 2D, E**). While the binding affinity of recombinant DVL2-FL (which contains all DVL protein domains) to cholesterol-PM vesicles is approximately 2-fold that of DVL2-PDZ, 7DHC-PM did not exhibit higher binding to DVL2-FL versus DVL2-PDZ (**Figure 2D-F**). As part the Wnt signalosome, DVL proteins form oligomers through interprotein swapping of the DEP (Beitia et al., 2021) or DIX domains (Kan et al., 2020) of DVL proteins. To determine if the increased PM binding of DVL2-FL compared to DVL2-PDZ in the presence of cholesterol but not 7DHC is due to DVL oligomer formation, PM binding was performed under non-denaturing conditions. Oligomer formation was only observed with DVL2-FL but not DVL2-PDZ (**Figure 2G**). Additionally, cholesterol-PMs promoted oligomerization more efficiently than 7DHC-PMs (**Figure 2G**). These data demonstrate that sterol structure is a determinant of DVL membrane association and suggest that sterol impaired oligomerization may affect downstream DVL signaling.

**Figure 2.**
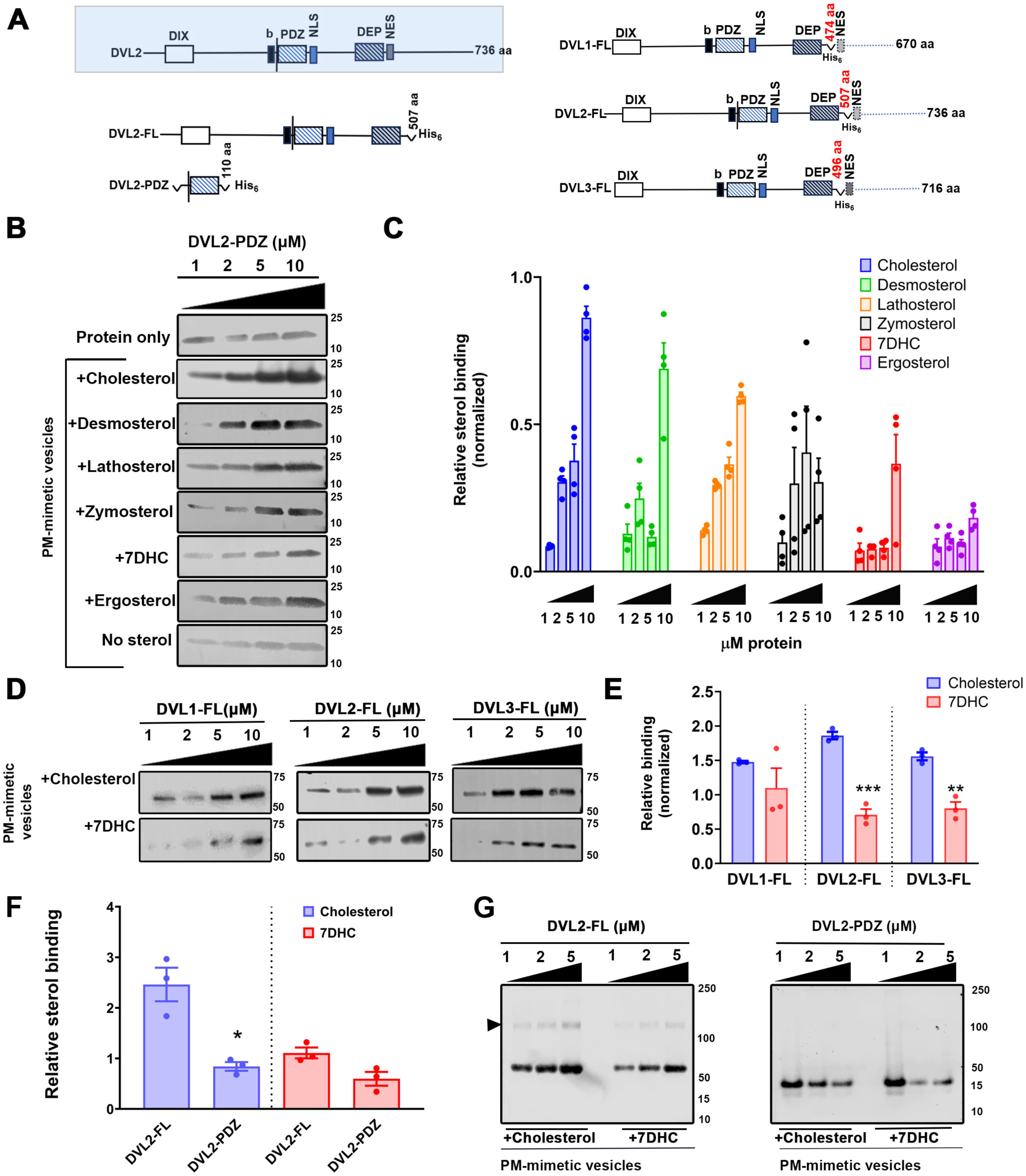
Sterol structure differentially impacts DVL membrane association. **A)** Domain structures of DVL isoforms and constructs utilized for this study. The critical functional domains, DIX, PDZ and DEP, in addition to basic amino acid-rich motif (b) and nuclear localization (NLS) and nuclear export (NES) signals, are shown. **B)** Representative western blots of PM-mimetic vesicle and DVL2-PDZ pulldowns. **C)** PM-mimetic vesicle and DVL2-PDZ pulldowns. N = 4 independent experiments. **D)** Western blot of DVL isoforms and PM-mimetic vesicle sedimentation assays. **E)** DVL isoform (10 µM) binding to PM-mimetic vesicle pulldown. N = 3 independent experiments. Data represent mean ± SEM; unpaired Student’s t-test. **F)** Quantified DVL2-FL and DVL2-PDZ (10 µM protein) from PM-mimetic vesicle sedimentation assays. N = 3 independent experiments. Data represent mean ± SEM; unpaired Student’s t-test. **G)** Coomassie staining of native PAGE of PM-mimetic vesicle pulldown assay, DVL2-FL vs DVL2-PDZ, with cholesterol and 7DHC containing PM-mimetic vesicles. *p < 0.05, **p < 0.01, ***p < 0.001, ****p < 0.0001 for all statistical tests.

### Sterol structure impacts the kinetics of DVL membrane association

To evaluate the kinetic parameters of DVL2-sterol binding, we analyzed the membrane affinity of DVL2-PDZ via surface plasmon resonance (SPR) analyses. Using liposome sensor chemistry with PM-mimetic vesicles captured on the hydrophobic sensor surface, we analyzed the binding of purified DVL protein to PM-mimetic vesicles. Combined comparative SPR sensorgrams for cholesterol versus alternate sterol binding to 40 nM recombinant DVL2-PDZ demonstrated sterol structure regulates the membrane binding efficiency of DVL2 (**Figure 3A**; **Figure S3**). K_D_ values demonstrate that while DVL2-PDZ shows a high affinity towards cholesterol, lathosterol, and desmosterol-rich PM-mimetic vesicles (K_D_ = 20 - 50 nM), DVL2-PDZ exhibited less affinity toward 7DHC, zymosterol, and ergosterol-rich PM-mimetic vesicles (K_D_ = 150 - 650 nM) (**Figure 3B**). Lipid-mediated membrane association of the DVL-PDZ domain was previously shown to require cholesterol to directly interact with the plasma membrane, with anionic lipids such as phosphatidylserine or phosphatidylinositol (PI) phosphates (PIPs) such as PI 4, 5-bisphosphate (PI(4,5)P_2_ or PIP2) also contributing (Sheng et al., 2014). Our SPR analyses determined that loss of either cholesterol or PIP2 caused a significant negative shift in 1:1 stochiometric DVL2 binding, suggesting that a localized and optimal concentration of cholesterol and PIP2 is imperative to increase the membrane affinity of the DVL2-PDZ (**Figure 3C**). While maximal B_max_ was achieved in the presence of 2% PIP2 and 8% cholesterol, a significant reduction was observed upon removal of either cholesterol or PIP2 (**Figure 3C**). Both PIP2 and cholesterol promoted DVL2 binding when added to POPC or POPC+POPS (**Figure 3C**). Increased binding for DVL2-FL was also observed compared to DVL2-PDZ in the presence of cholesterol **(Figure 3E, 3F).** While K_D_ values demonstrated cholesterol differentially impacted DVL2-FL versus DVL2-PDZ binding (**Figure 3G**), the presence of 7DHC had no impact on DVL2-FL versus DVL2-PDZ binding. These data demonstrate sterol structure promotes optimal DVL protein membrane association, likely in cooperation with localized PIP protein association.

**Figure 3.**
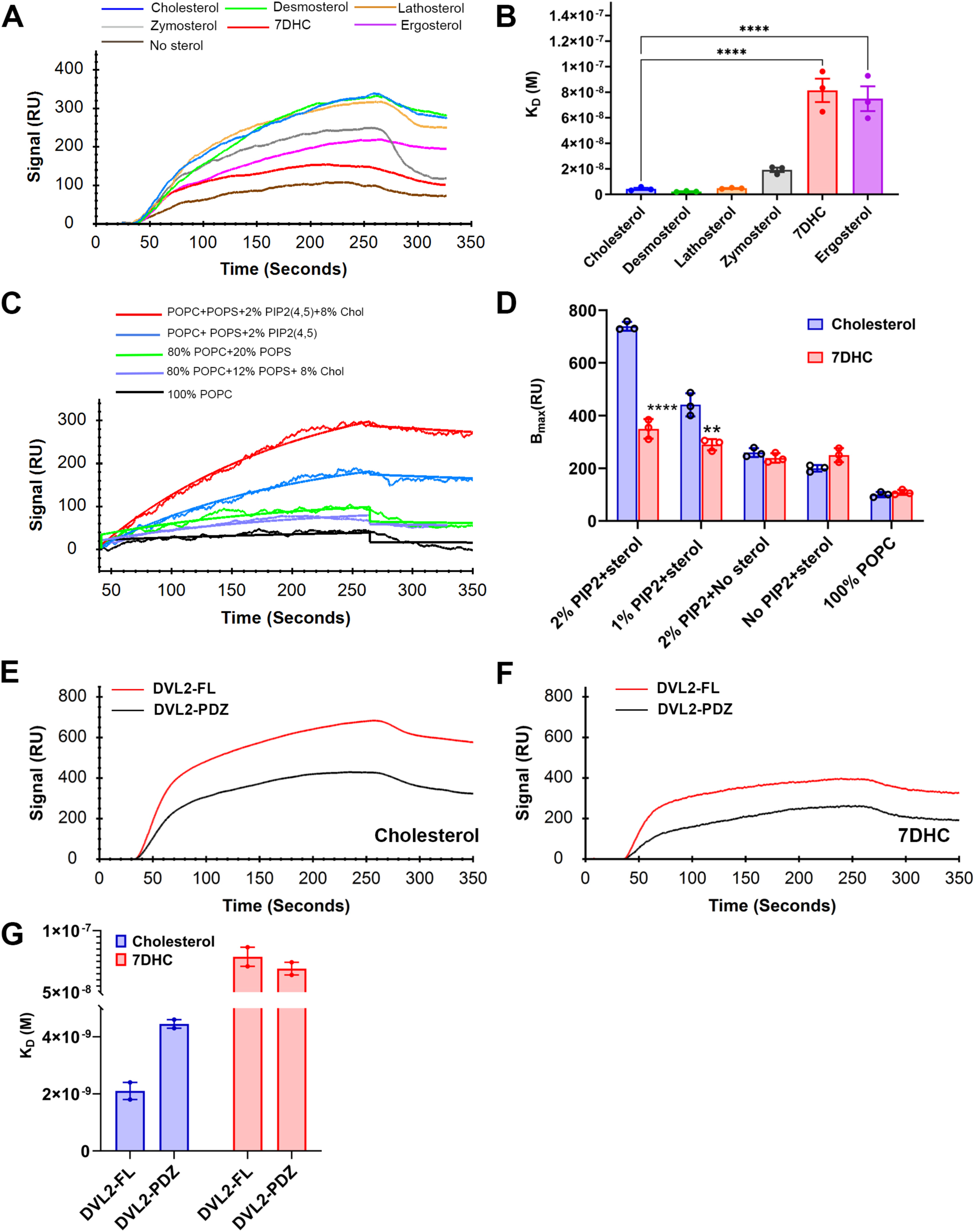
Sterol structure regulates the kinetics of DVL membrane association. **A)** Representative binding curves (combined sensorgrams) of DVL2-PDZ domain with sterol-incorporated PM-mimetic vesicles. **B)** K_D_ values of DVL2-PDZ domain binding with sterol-incorporated PM-mimetic vesicles. N = 3 independent experiments. Data represent mean ± SEM. One-way- ANOVA with Brown-Forsythe test and Dunnet’s test. **C)** Representative SPR binding curves (combined sensorgrams) showing phosphatidylinositol (4,5) bisphosphate (PIP2) dependence of DVL2-PDZ domain for PM-mimetic vesicle binding. **D)** Maximum binding signal for DVL2-PDZ for PM-mimetic vesicles in the presence or absence of PIP2. N = 3 independent experiments. Data represent mean ± SEM. Unpaired student’s t-test. **E**) Representative binding curves (combined sensorgrams) for DVL2-FL (red curve) versus DVL2-PDZ (black curve) to PM-cholesterol; protein concentration = 40 nM. **F)** Representative binding curves for DVL2-FL (red) versus DVL2-PDZ (black) to PM-7DHC; protein concentration = 40 nM. **G)** K_D_ values of DVL2-FL and DVL2-PDZ to PM-cholesterol and PM-7DHC. N = 2 independent experiments. Data represent mean ± SEM. **p < 0.01, ****p < 0.0001 for all statistical tests.

### Disrupted sterol homeostasis shifts DVL2 localization from the plasma membrane to the nucleus

Though it is predominantly found cytosolically and much work has focused on the inner plasma membrane association of DVLs with receptors such as FZD7 and LRP5/6, DVLs are also expressed within the nucleus (Castro-Piedras et al., 2021; Itoh et al., 2005). NES and NLS sequence motifs allow DVL isoforms to localize to the nucleus where DVLs localize with various transcription factors and cell division machinery (Itoh et al., 2005). Although the nuclear role of DVLs are still being elucidated, nuclear DVLs are reported to bind promoters such as CYP19A1 in breast cancer cell lines (Sharma et al., 2021a; Sharma et al., 2019) and to regulatory proteins in Wnt responsive gene transcription such as c-Jun and c-Myc (Gan et al., 2008). DVL also binds to chromatin modifiers such as KM2D, and EZH2 in breast cancer cells (Castro-Piedras et al., 2021; Sharma et al., 2021b). We hypothesized that given the requirement of cholesterol for DVL2 membrane association, cholesterol depletion would inhibit this process, augmenting DVL2 subcellular localization and function in other cellular compartments. To test this, HEK293T cells were grown in cholesterol-depleted media (LPDS) and treated with AY9944 to inhibit DHCR7- mediated cholesterol synthesis **(Figure 4A)**. This treatment reduces overall cellular sterols, reduces cholesterol content, and causes accumulation of 7DHC (Anderson et al., 2021)(**Figure S4A-D)**. To determine whether changes in DVL2 compartmentalization are due to cholesterol depletion or the effect of specific accumulating sterols, cells were treated in parallel with the HMG-CoA reductase inhibitors Simvastatin or Atorvastatin, or the DHCR24 inhibitor U18666A (**Figure 4A**). GC-MS analyses of sterol content showed reduction of cholesterol and accumulation of appropriate alternate sterols under respective treatments (**Figure S4D**). While total cellular DVL2 accumulation was unchanged after cholesterol synthesis inhibition, indicating cholesterol depletion does not significantly affect DVL2 turnover (**Figure 4B, C**), we observed that inhibitor treatments increased nuclear accumulation of DVL2 at the expense of membrane association (**Figure 4D, E**). Increased nuclear localization of DVL2 was further confirmed using confocal imaging of a previously validated CRISPR-Cas9 gene edited HEK293T cell line with DVL2 endogenously tagged with a C-terminal mEGFP (Ma et al., 2020)(**Figure 4F**). Both AY9944 and Simvastatin treatment induced mEGFP accumulation within the nucleus to varying degrees (**Figure 4F, G, H**), suggesting overall sterol levels and sterol structure may both impact DVL localization. These data demonstrate a previously unrecognized impact of altered sterol content on induction of DVL nuclear localization.

**Figure 4.**
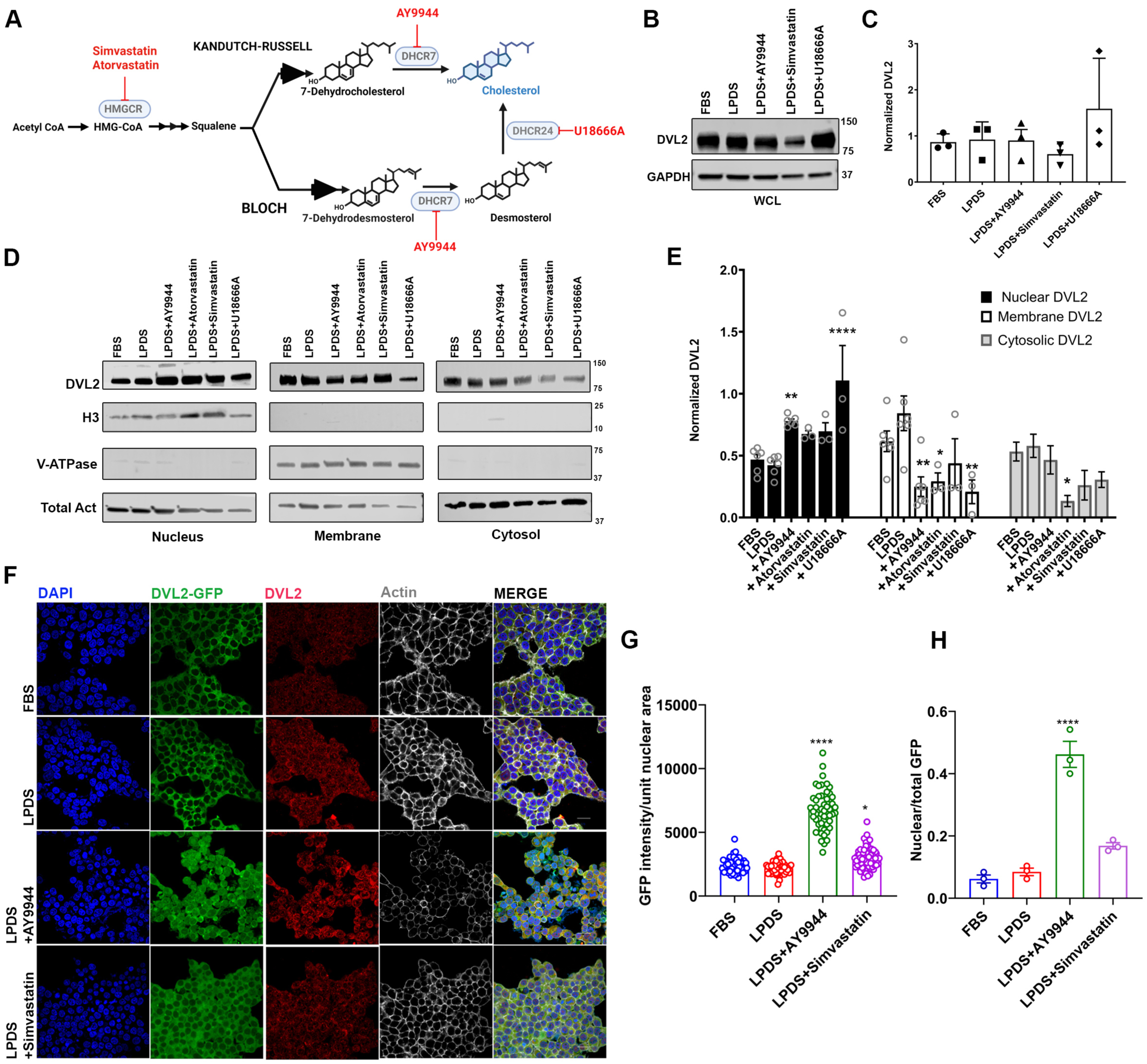
DVL2 subcellular localization is shifted in response to sterol change. **A)** Schematic illustrating the final steps of post-squalene cholesterol synthesis. Small molecule inhibitors are indicated in red, target proteins in blue. **B)** Western blot of total DVL2 expression following inhibitor treatments. **C)** Total cellular DVL2 normalized to GAPDH. N = 3 independent experiments. Data represent mean ± SEM. One-way ANOVA with Dunnett’s test. **D**) Representative western blots of subcellular DVL2 following inhibitor treatments. **E)** Subcellular DVL2 normalized to total actin. N = 3 independent experiments. Data represent mean ± SEM. One-way ANOVA with Dunnett’s test. **F**) Confocal imaging of HEK293T-DVL2-mEGFP-KI cells following respective treatments**. G)** Nuclear GFP/unit nuclear area. N = 48-55 individual cells ± SEM. One-way ANOVA (F (3, 218) = 274.3, P<0.000001) and Dunnett’s test. **H)** Nuclear GFP normalized to total GFP/field. N = 3 independent experiments. One-way ANOVA (F (3, 8) = 62.68, P<0.0001) and Dunnett’s test. *p < 0.05, **p < 0.01, ***p < 0.001, ****p < 0.0001 for all statistical tests. Scale bar = 25 µm.

### Nuclear localization of DVL2 is dependent on PDZ-domain interaction with FoxK2

It has previously been reported that nuclear translocation of DVL2 requires FoxK2 binding with the PDZ domain of DVL2 (Wang et al., 2015). Upon cholesterol synthesis inhibition with AY9944, co-immunoprecipitation (Co-IP) demonstrates nuclear FoxK2-DVL2 association is increased **(Figure 5A and B)**. We confirmed that FoxK2 expression itself was not affected by cholesterol synthesis inhibition (**Figure S5A and B**). To determine if FoxK2 binding to the DVL2-PDZ domain is required for DVL2 nuclear localization in our model, we utilized an inhibitor of the DVL2-PDZ domain, NSC668036 (Shan et al., 2005). The predicted binding site of NSC668036 to the DVL2-PDZ domain overlaps with the predicted cholesterol binding pocket of DVL2-PDZ and major FoxK2 binding residues within the DVL2-PDZ domain (**Figure 5C**). While treatment of cells with NSC668036 in cholesterol-rich conditions did not affect DVL2 nuclear expression, NSC668036 treatment inhibited binding of DVL2 and FoxK2 within the nucleus (**Figure 5G**) and completely antagonized AY9944-mediated nuclear shuttling **(Figure 5D-F)**, further suggesting DVL2-Foxk2 association occurs upon cholesterol loss. These data demonstrate nuclear localization of DVL2 within cholesterol synthesis inhibited conditions is FoxK2-dependent.

**Figure 5.**
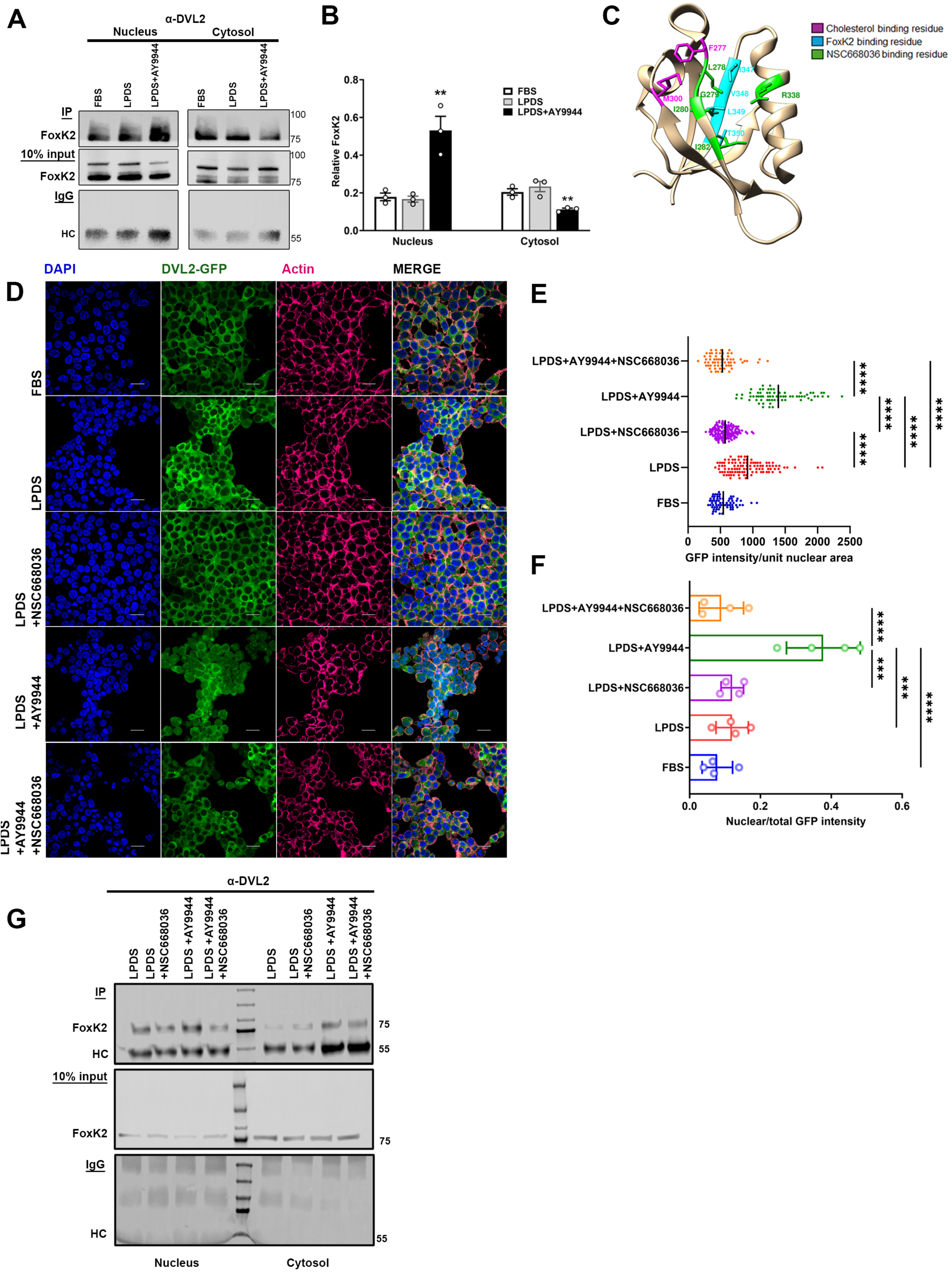
Nuclear DVL2 transport is FoxK2 dependent and sensitive to PDZ-domain inhibition. **A)** Co-IP between DVL2 and FoxK2 under sterol-impacted conditions. **B)** Nuclear DVL2 binding to FoxK2. N = 3 independent experiments. Data represent mean ± SEM. One-way ANOVA with Dunnett’s test; F (2, 6) = 20.58, P=0.0021 (nucleus); F (2, 6) – 11.22, P=0.0094 (cytosol). **C)** Model for NSC668036, FoxK2, and cholesterol binding to DVL2-PDZ. Predicted NSC668036 binding residues are green, cholesterol residues are purple, FoxK2 residues are cyan. **D**) HEK293T-DVL2-mEGFP-KI cells following sterol treatments. **E)** Nuclear GFP/unit nuclear area. N > 50 cells from 2 independent experiments. Data represent mean ± SEM. One-way ANOVA (F (4, 438) = 168.5,P<0.0001) with Tukey’s test. **F)** Nuclear GFP normalized to total GFP/field. N = 4 fields from 2 independent experiments. One-way ANOVA (F (4, 15) = 15.74, P<0.0001) and Tukey’s test. **G)** Co-IP between DVL2 and FoxK2. *p < 0.05, **p < 0.01, ***p < 0.001, ***p < 0.0001 for all statistical tests. Scale bar = 25 µm.

### Loss of sterol homeostasis impacts DVL2 protein-protein interactions

To determine how changes in sterol content regulate DVL2 protein binding partners and cell signaling, DVL2 was tagged with the promiscuous biotin ligase TurboID to biotinylate DVL2 interacting proteins (Branon et al., 2018; May et al., 2020). Through analyses of DVL2 interacting partners in HEK293T cells stably expressing DVL2-TurboID fusion protein and cultured in FBS, LPDS, or LPDS with AY9944, we identified a total of 381 candidate DVL2 interacting partners in FBS conditions, 127 DVL2 interacting proteins in LPDS conditions, and 124 DVL2 interacting candidates in AY9944-treated cells (**Figure 6A; Table S5**). A total of 88 protein interactors were present in all three conditions, suggesting those interactions are likely not regulated by sterol availability. Of the condition exclusive protein interactions, 240 interactions were specific to FBS, 7 interactions were specific to LPDS, and 12 interactions were specific to AY9944 treated cells (**Figure 6A**). Predictive analysis in STRING database of the 240 interactions lost in FBS culture include important DVL-associated networks including clathrin-mediated endocytosis, basal body anchoring, and vesicle-mediated transport (**Figure 6B**). DVL2 interaction networks which were not lost by sterol changes also exhibited functional change. For example, the DVL2/tumor suppressor Tp53 (Lim et al., 2006; Rual et al., 2005) network was maintained following AY9944 treatment but exhibited reduced complexity (**Figure 6C**) with increased DVL2-Tp53 association **(Figure 6D**). By integrating Tp53 interactors within the STRING database, known Tp53 target proteins were also identified as possible DVL2 interactors (**Figure 6C**). DVL2 interactions with known binding partners β-catenin and Axin1/2 were also maintained (**Figure 6F**) but DVL2 association was altered. Co-IP between DVL2 and β-catenin showed increased binding within the nucleus in cholesterol-depleted conditions **(Figure 6G)**. Axin2 exhibited decreased interaction within both the cytosol and nucleus following AY9944 treatment (**Figure 6H**). Overall, these data demonstrate loss of sterol homeostasis dramatically impacts DVL2 protein interaction networks with potentially significant impacts on cell signaling and function.

**Figure 6.**
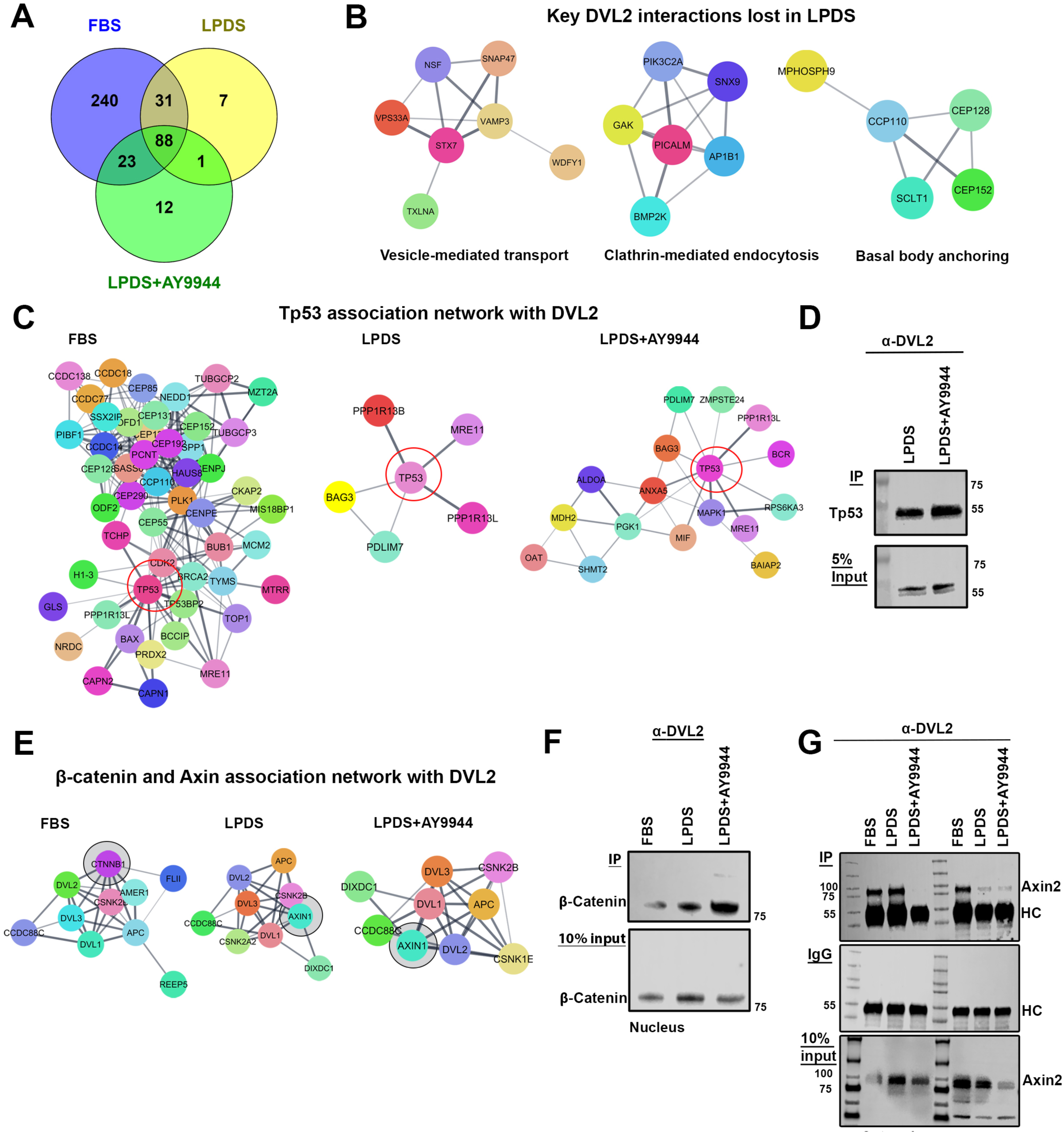
Loss of sterol homeostasis alters the DVL2 protein-protein interaction network. **A)** Venn diagram of DVL2-interacting proteins in HEK293T cells under sterol-specific conditions. **B)** Representative key DVL2 interactions lost within LPDS treatment. **C)** DVL2-Tp53 interaction network in FBS, LPDS, and AY9944 treatment. **D)** Co-IP between endogenous DVL2 and p53 in HEK293T. **E)** β-catenin and Axin1/2 interaction networks with DVL2 under sterol-specific conditions. **F)** β-catenin and DVL2 interaction within the nucleus. **G)** Co-IP of endogenous DVL2 and Axin2 in HEK293T cells.

### Nuclear expression of DVL2 is increased within developmental models of cholesterol synthesis disorders

To determine if increased nuclear DVL2 is also found within developmental and disease models associated with sterol changes, we analyzed DVL2 in human neural stem cells (NSCs) derived from induced pluripotent stem cell (iPSC) models of SLOS (Francis et al., 2016). While expression of neither total nor cytosolic DVL2 was altered in SLOS NSCs (**Figure 7A, B**), we found increased nuclear DVL2 in SLOS NSCs or AY9944-treated control NSCs (**Figure 7A, B**). Membrane-localized expression of DVL2 was stabilized by cholesterol supplementation (**Figure 7A, B**). We also analyzed DVL2 localization in iPSCs following acute removal of cholesterol by methyl-β-cyclodextrin (MβCD), AY9944, or Simvastatin treatment. While both AY9944 and simvastatin altered DVL2 localization following cholesterol depletion, acute stripping of membranous cholesterol by MβCD did not affect DVL2 localization (**Figure 7C, D**). To demonstrate nuclear accumulation of DVL2 occurs *in vivo*, we analyzed DVL2 expression within a mouse model of SLOS (*Dhcr7* ^Δ3-5^/^T93M^) (Correa-Cerro et al., 2006; Freel et al., 2022; Wassif et al., 2001). Analyses of DVL2 within isolated cerebral cortices showed a significant increase in nuclear DVL2 and reduction in membrane-associated DVL2 in *Dhcr7* ^Δ3-5^/^T93M^ mice compared to *Dhcr7^+/^*^T93M^ controls (**Figure 7E, F**). Overall, these data demonstrate that nuclear accumulation of DVL2 in response to altered sterol homeostasis is a physiological effect observed in cellular and animal models of human disease.

**Figure 7.**
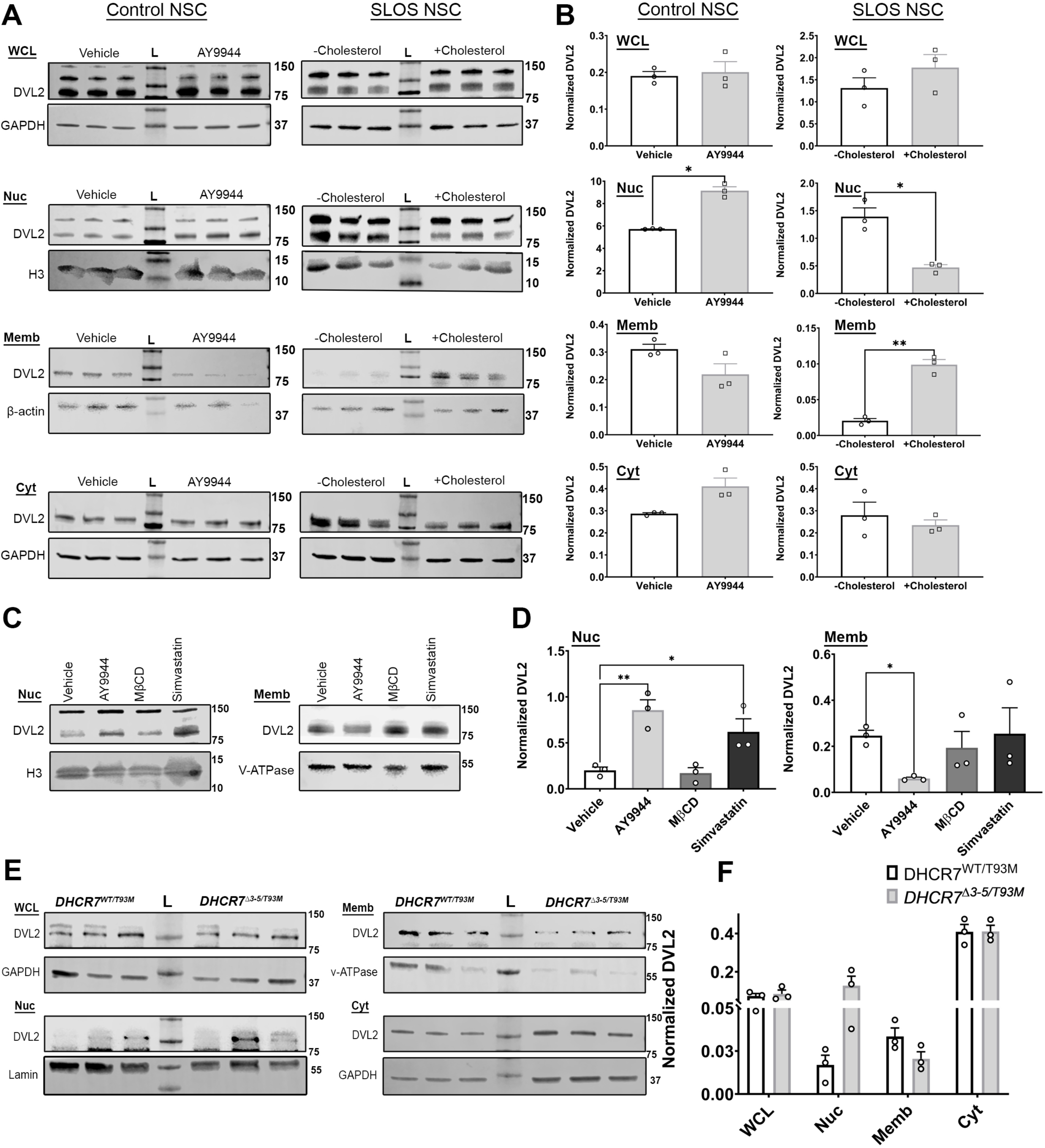
Nuclear translocation of DVL2 in genetic and pharmacological models of Smith-Lemli-Opitz syndrome. **A**) Representative western blots of DVL2 in subcellular fractions of NSCs derived from control and SLOS iPSCs. **B**) DVL2 accumulation in SLOS and control NSCs. N = 3 independent experiments. Data represent mean ± SEM. Unpaired Student’s t test with Welch’s correction. **C**) Representative western blots of endogenous DVL2 accumulation in subcellular fractions of control iPSCs. **D**) DVL2 expression in fractionated iPSCs. N = 3 independent experiments. Data represent mean ± SEM. One way ANOVA with Dunnett’s test for nucleus (F (3, 8) = 11.87, P=0.0026) and membrane (F (3, 8) = 1.765, P=0.2314). **E**) Representative western blots for DVL2 expression in cerebral cortices of *Dhcr7*^Δ3-5^/^T93M^ and *Dhcr7*^WT^/^T93M^ mice. **F**) DVL2 expression in cerebral cortices. N = 3 independent experiments. Data represent mean ± SEM. Unpaired Student’s t test with Welch’s correction. * p<0.0001 * p < 0.05, ** p < 0.01, and *** p < 0.001 for all statistical tests. WCL = whole cell protein lysate; Nuc = nuclear protein fraction; Cyt = cytosolic protein fraction; Memb = membrane protein fraction; L = protein ladder.

## DISCUSSION

Herein, we report that structurally distinct sterols of clinical and biological significance are detrimental to normal DVL protein localization and function, resulting in aberrant nuclear localization and disruption of DVL protein-protein interactions (**Figure 8**). The structure of cholesterol allows it to pack tightly with other membrane lipids and integral proteins. The α-face allows interaction with saturated fatty acyl chains of phospholipids or sphingolipids or stacking against side chains of aromatic amino acids; the β-face interacts with projected methyl groups of unsaturated fatty acids or branched amino acids of transmembrane protein domains (Lange and Steck, 2008). The heterogeneous distribution of cholesterol between the membrane leaflets and the higher degree of fatty acid unsaturation in the inner plasma membrane makes the inner membrane less rigid and more conducive to lipid diffusion (Ingólfsson et al., 2014). While the exact ratio of inner to outer plasma membrane sterols remains controversial (Buwaneka et al., 2021; Courtney et al., 2018; Liu et al., 2017; Pham et al., 2022), it is plausible that sterol flipping contributes to locally enriched cholesterol within the inner membrane to regulate DVL signaling (Sheng et al., 2014).

**Figure 8.**
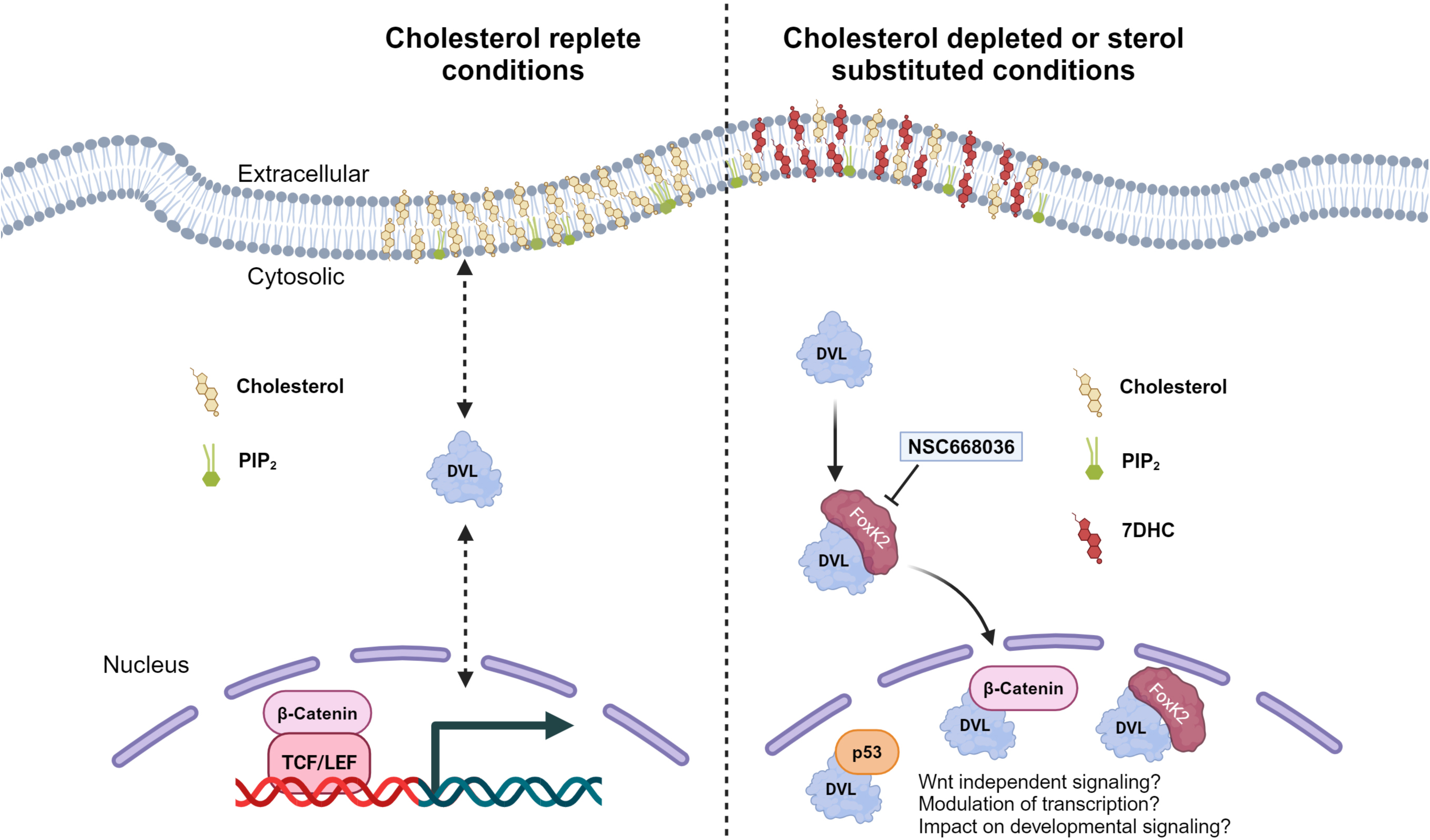
Sterol-dependent DVL subcellular localization and potential cellular impacts. Maintenance of local cholesterol abundance and PIP2 concentrations allow DVL2 to shuttle between intracellular compartments to bind to the inner plasma membrane. Upon cholesterol loss or changes to the sterol composition of the plasma membrane, DVL2 plasma membrane association is impaired and results in nuclear localization of DVL2. The multifaceted scaffolding properties of DVL may impact a variety of cellular processes important for cell and tissue function.

Our data also suggests that DVL membrane association is likely co-dependent on sterol and PIP2 protein association. Removal of either cholesterol or PIP2 significantly reduced the binding of the DVL-PDZ domain to the membrane (**Figure 3**). Due to the absence of a cholesterol-binding motif within the DVL2-PDZ domain, the role of neighboring membrane lipids in mediating DVL membrane recruitment is plausible. It has been recently reported that DVL3 is required to organize PI4K and PIP5K in coordination to synthesize PIP2 (de la Cruz et al., 2022). The ability of a sterol to compensate for cholesterol in a stable phospholipid background is likely dependent on the ability of that story to support L_o_ of lipids within the membrane. Model membranes or cellular models that incorporate 7DHC exhibit high fluidity and altered phospholipid packing (Chen and Tripp, 2012; Tulenko et al., 2006). AY9944-treated rats exhibited differential protein expression in L_o_ lipid rafts, suggesting that sterol composition impacts protein localization at the membrane (Keller et al., 2004). However, sterol influence on lipid packing is likely sterol specific (Huster et al., 2005). While 7DHC increases lipid movement within the fluid phase of membranes, cholesterol and desmosterol reduce the free volume (Shrivastava et al., 2008). Therefore, the impact of sterol biochemistry on liquid ordering and subsequently lipid accessibility could also play an important role in DVL-membrane association.

Dishevelled has historically been categorized as a modulator of Wnt/β-catenin signaling, acting as a balance between canonical, β-catenin-dependent transcriptional programs and non-canonical, Ca^2+/^polar cell polarity signaling (Gordon and Nusse, 2006; Schlessinger et al., 2007). Wnt-independent roles of DVL proteins have been largely understudied. We observed that reduction of membrane associated DVL2 in response to changes in sterol content is accompanied with increased nuclear DVL2 (**Figure 4D-H**) independent of changes in total DVL2 (**Figure 4B**), suggesting translocation of cytoplasmic DVL2 to nucleus. While the exact functions of nuclear DVL require elucidation, several studies have suggested DVL proteins can function as transcriptional regulators. For example, nuclear DVL2 promotes transcriptional activation of Wnt target genes via c-Jun association to stabilize β-catenin/TCF interactions (Gan et al., 2008). DVL proteins have also been reported to be recruited to promoter sequences to complex with β-catenin and LEF1/TCF4 to positively modulate canonical Wnt (Gan et al., 2008; Torres and Nelson, 2000). Nuclear DVL proteins also exhibit Wnt-independent functions. ChIP sequencing demonstrated strong association of DVL1 and DVL3 with promoters for genes that regulate the cell cycle, chromatin organization, DNA repair, signal transduction, and immune system modulation (Boligala et al., 2022; Castro-Piedras et al., 2021; Rasha et al., 2023). Nuclear DVL1 and DVL3 were also essential for myoblast proliferation in a β-catenin-independent manner (Pruller et al., 2022). DVL proteins may also impact the epigenome, acting as a scaffold for the epigenetic modifier lysine methyltransferase 2D (KMT2D) (Castro-Piedras et al., 2021). Whether nuclear accumulation of DVL in response to changes in sterol biochemistry exerts similar cellular impacts requires further study. Further analyses of sterol impacts on DVL localization and function under both Wnt stimulated versus standard conditions is required.

The signaling required and molecular partners which mediate DVL localization in response to sterol changes are unclear. Forkhead box transcription factors, such as FoxK1 and FoxK2, increase Wnt/β-catenin signaling by translocating DVL into the nucleus (Wang et al., 2015). However, additional mechanisms may be at play. Nuclear shuttling of DVL monomers is partially regulated by two conserved NES and NLS sequences (Itoh et al., 2005; Sokol, 1996). Also, supramolecular intracellular condensates of DVL2 colocalize with centrosomal proteins such as γ-globulin and CEP164 in a Wnt-dependent and cell cycle-dependent manner (Schubert et al., 2022). Lastly, post-translational modification of DVL residues through acetylation or phosphorylation can also regulate DVL nuclear accumulation (Sharma et al., 2019), as well as DVL protein-protein interactions through PDZ domain activity (Jurasek et al., 2021). A previous study linked DVL3 phosphorylation patterns to its function (Hanáková et al., 2019). Differential phosphorylation of DVL in response to sterol biochemistry could signal for nuclear import. Dissection of these potential mechanisms for increased DVL nuclear translocation following sterol change are required.

Our work also demonstrates that sterol-associated changes in DVL localization impair DVL protein-protein interactions. We observed a significant attenuation of DVL2 protein-protein interactions upon sterol depleted culture, including DVL2-protein interactions involved in clathrin-mediated endocytosis, ER to Golgi anterograde transport, centrosome function, cellular trafficking, and transcriptional regulation by β-catenin and Tp53 (**Figure 6; Table S5**). Sterol regulated association of Tp53 with DVL2 may play a role in Tp53 tumor suppressor activity. It has been shown that silencing of the *Wwox* tumor suppressor increases nuclear DVL2 in head and neck cancer (Celebi et al., 2020). We also observed significant overlap between DVL2 and p53 interacting protein networks (**Figure 6C**). These data suggest DVL protein-protein interactions are likely exquisitely sensitive to sterol content. Loss of protein interactions indicates changes in protein partners between culture conditions and possibly intercompartmental shift of protein networks.

The regulation of DVL by membrane lipid content has significant connotations for cell specific and disease-associated impacts of DVL. For example, individuals with Robinow syndrome caused by autosomal-dominant mutations within *DVL1* or *DVL3* display tissue specific phenotypes associated with skeletal malformations, genital abnormalities, and cognitive dysfunction (Schwartz et al., 2021; White et al., 2016) while other tissues are largely spared. Similar tissue specific findings have also been observed in mouse, chicken, and drosophila models, indicating a conserved tissue response across species (Gignac et al., 2023). Due to the functional redundancy of DVL proteins (Lee et al., 2008), a molecular regulator such as sterol availability which can impact all DVL isoforms simultaneously may exacerbate DVL tissue defects. Additionally, sterol content and expression vary both across cell types and temporally. For example, on-tissue based liquid chromatography/mass spectrometry of the brain revealed dramatically different expression profiles of sterols depending on brain region and cell type (Allen et al., 2019; Yutuc et al., 2020). There is also a well-established association of altered sterol homeostasis with aging and neurodegeneration (Essayan-Perez and Sudhof, 2023; van der Kant et al., 2019; Varma et al., 2021). Given the known requirements of DVL and Wnt/β- catenin signaling during neurodevelopment (Rosso et al., 2005; Zheng et al., 2015), further defining sterol specific impacts on DVL signaling could have broad scientific and clinical significance. Utilizing cellular or animal models which would allow analyses of temporal and cell specific impacts of sterol biochemical change on DVL activity will be essential to defining the biological importance of this critical signaling molecule.

This work may also be important in defining the biological impacts of sterols within sterol-impacted rare diseases. Defects in cholesterol biosynthesis are responsible for eight known rare disorders due to mutations within post-squalene enzymes which catalyze steps of the Kandutsch-Russell or Bloch cholesterol synthesis pathways (Porter and Herman, 2011). Despite patients sharing a cholesterol deficient biochemistry, these syndromes exhibit distinct clinical phenotypes (Porter, 2002). One hypothesis is the structural dissimilarity among accumulating sterols may disparately impact sterol-sensitive signaling pathways. Our previous work reported 7DHC accumulation inhibited Wnt /β-catenin signaling through impaired DVL2 activity (Francis et al., 2016). However, we also showed in this same study that Wnt signaling was not inhibited by lathosterol accumulation (Francis et al., 2016). This current study now confirms our previous cellular observation at the molecular level, demonstrating that clinically relevant sterols such as lathosterol can compensate for cholesterol regarding DVL membrane affinity (**Figure 4**). These data suggest that sterol-specific impacts on the function of DVLs or other sterol-interacting proteins play a role in the clinical phenotypes observed in cholesterol synthesis disorders (Francis et al., 2016).

In summary, this study demonstrates that DVL membrane association is regulated by membrane sterol content with significant impacts on DVL localization and function. The results of our binding studies were obtained in the absence of exogenous Wnt ligand stimulation, which suggests that the PDZ domain has intrinsic membrane binding properties defined by sterol-phospholipid signatures in the membrane. The dual role of PDZ domain in both membrane and nuclear localization is notable and indicates that the scaffolding and protein-protein interaction properties of DVL at membrane organelles may be important in other signaling pathways. Our work demonstrates sterols exert broad impacts on DVL activity and localization, possibly through both altered lipid organization and DVL protein-protein interactions. Additional studies are required to fully delineate the impact of DVL-sterol interactions on mammalian health and disease.

## Supporting information

Supplemental Information_Sengupta et al

Table S5_Sengupta et al.

## ACKNOWLEDGEMENTS

This study was supported by the National Institutes of Health (NIGMS P20GM103620, P20GM103548). M.M.S. was supported by the Sanford Program for Undergraduate Research (P20 GM103620). We would like to thank Anna Sundborger (Uppsala University) for assistance with preparation of PM-mimetic vesicles. We would also like to thank Wonhwa Cho (University of Illinois at Chicago) for sharing plasmids and scientific discussion. Thank you to Xi He (Harvard University) for sharing of HEK293T DVL2-mEGFP-KI cells. Thank you to Alex Sodt (*Eunice Kennedy Shriver* National Institute of Child Health and Human Development) for helpful discussions. We would like to thank the Center for Brain and Behavior Research at the University of South Dakota for supporting this project. We would like to thank the following core facilities at Sanford Research for experimental support: Imaging, Biochemistry, and Functional Genomics and Bioinformatics cores. We would also like to thank the Proteomics Shared Resource Facility at Sanford-Burnham-Prebys for proteomics services. Illustrations were created using BioRender (https://biorender.com/). Any opinions, findings and conclusions expressed in this material are those of the author(s) and do not necessarily reflect the views of the National Institutes of Health.

## AUTHOR CONTRIBUTIONS

Conceptualization, S.S. and K.R.F.; methodology, S.S., J.D.W.Y., M.M.S., and K.R.F.; investigation, S.S., J.D.W.Y., and M.M.S.; writing—original draft, S.S., K.R.F.; writing—review & editing, S.S., J.D.W.Y., M.M.S., and K.R.F.; funding acquisition, K.R.F.; supervision, K.R.F.

## MATERIALS AND METHODS

### Homology modeling and docking

To build a model of the DVL PDZ domain-cholesterol complex, the crystal structure of the DVL2 PDZ Domain in complex with the N2 Inhibitory Peptide (PDB ID: 3CBZ) was used. A cholesterol molecule was docked first onto the free PDZ domain structure using Autodock Vina application (Trott and Olson, 2010) in the UCSF Chimera platform (Pettersen et al., 2004). Sterol three-dimensional conformers were obtained from Pubchem and docked on the obtained crystal structure of DVL2-PDZ domain as described (Trott and Olson, 2010).

### Protein expression and purification

*Dvl2*-PDZ-pET-30a (+) bacterial expression construct was a kind gift from Wonhwa Cho (University of Illinois, Chicago). The complimentary DNAs (cDNAs) for other domains of human *Dvl1*, *Dvl2* and *Dvl3* were cloned into pET-30a (+) (Invitrogen) vector using In-Fusion® HD Cloning Kit (Takara Bio USA, Inc). Primer sequences used were as follows: *hsDvl1* (forward; 5’- AGATATATATACATATGATGGCGGAGACCAAGATTATC-3’, reverse; 5’- GGTGGTGGTGCTCGAGGAGATCCCCGAAGACGTAG-3’) *hsDvl2* (forward; 5’- AGATATATATACATATGATGGCGGAGACCAAGATTATC-3’, reverse; 5’- GGTGGTGGTGCTCGAGTCTCCGAAGACGTAATAGCA-3’) *hsDvl3* (forward; 5’- AGATATATATACATATGATGGGCGAGACCAAGATCATC-3’, reverse; 5’-GGTGGTGGTGCTCGAGGTCACCGAAGATGTAGTAGCACTG-3’). All recombinant proteins were expressed with an C-terminal His-tag using pET-30a vector. All expression constructs were transformed into BL21-CodonPlus (DE3)-RIL cells (Agilent Technologies, 230245) for protein expression. A 10 mL preculture was established overnight from a single transformed colony in LB medium supplemented with 50 µg/mL Kanamycin (Sigma Aldrich, PHR1487).

Cells were induced with 1 mM Isopropyl ß-D-1-thiogalactopyranoside (IPTG, Sigma Aldrich, I6758) for 4 h at 25° C. Expressed proteins were purified to homogeneity using a 1 mL His-Trap column (Cytiva). During purification, 2 mL fractions were collected in stepwise gradient of imidazole (Sigma-Aldrich, 15513). The purity of isolated fractions was validated by 4-10% SDS-PAGE (BioRad) (**Figure S2A-C)**. Protein fractions were then desalted and concentrated using Pierce concentrators with a 3 kDa cut-off. Protein was further validated via western blot with an anti-His antibody (Cell Signaling, 2365T). Protein concentration was determined by BCA assay (Fisher Scientific, PI23227), aliquoted in 20 mM Tris-HCL, 100 mM NaCl, 5% glycerol, and stored at −80° C.

### Preparation of plasma membrane-mimetic vesicles

Plasma membrane (PM)-mimetic vesicles were prepared following previously described inner plasma membrane lipid composition with slight modifications (Sheng et al., 2014). All lipids were purchased from Avanti lipids or Cayman Chemicals. Lipid stock solutions were dissolved in chloroform (10 mg/mL). Briefly, a mixture of 1-palmitoyl-2-oleoyl-sn-glycero-3-phosphocholine (POPC, Avanti Polar Lipids, 850457), 1-palmitoyl-2-oleoyl-snglycero-3 phosphoethanolamine (Avanti Polar Lipids, POPE, 850757), 1-palmitoyl-2-oleoyl-sn-glycero-3-phosphoserine (Avanti Polar Lipids, POPS, 840034), cholesterol, 1-palmitoyl-2-oleoyl-sn-glycero-3-phosphoinositol (Avanti Polar Lipids POPI, 850142), and 1,2-dipalmitoyl derivatives of phosphatidylinositol-(4,5)-bisphosphate (PtdIns(4,5)P_2_, Cayman Chemicals, 64924) was prepared in a molar ratio of 12:35:22:22:8:1 in glass vials for pulldown assays. The lipid mixture was dried into a thin film under nitrogen gas and desiccated overnight at room temperature to remove organic traces.

Lipids were rehydrated in either HKM buffer (20 mM HEPES pH 7.4, 150 mM potassium acetate, 1 mM magnesium chloride, for pulldown assay) or in HBS buffer (20 mM HEPES, 150 mM NaCl, pH 7.4, for SPR assay) to a working concentration of 10 mM total lipid. To achieve homogenous 100 nm vesicles, the lipid mixture was freeze-thawed and sonicated, followed by sequential extrusion through polycarbonate membranes of appropriate pore sizes (800 nm, 400 nm, 100 nm; Avanti Polar lipids, 61009, 61007, 61010 respectively) via a mini extruder (Avanti Polar Lipids, 610020).

For optimization of sterol concentration in PM-mimetic vesicles in SPR assays, a 5-22% molar concentration of sterol was analyzed for optimal binding with DVL2-PDZ. The final working concentration of sterol for comparative SPR assays was chosen to be 8% based on optimal difference in relative unit (RU) between different sterols. The homogeneity of PM-mimetic vesicle size was validated via nanoparticle analysis (NanoSight NS300, Malvern Panalytical). PM-mimetic vesicles were diluted 1:6 in PBS and particle size obtained via diffraction analysis (**Figure S2D-F**). Sterol incorporation into PM-mimetic vesicle was quantitatively validated by GC-MS (**Figure S2G-H).** Sterol content within PM-mimetic vesicles was normalized to the internal standard coprostanol.

### PM-mimetic vesicle-protein sedimentation assay

Purified protein was added to PM-mimetic vesicles at varying concentrations as noted per experiment. The reaction was incubated at room temperature for 60 min. Protein bound to the PM-mimetic vesicle fraction was obtained by ultracentrifugation at 50,000 rpm at 4° C. The pellet fraction containing lipid-bound protein was isolated from unbound protein within the supernatant. Lipid-bound protein was run on 4-20% SDS-PAGE gel and immunoblotted with His-tag antibody (rabbit monoclonal, Cell Signaling Technology, 2365). Blots were visualized on a Li-Cor Odyssey imaging system. Protein quantification was normalized relative to sterol-free PM-mimetic vesicles.

### Surface plasmon resonance assay

Surface plasmon resonance (SPR) measurements were obtained using a one channel OpenSPR system (Nicoya). Assays were performed at 20 μL/min injection speed in HBS running buffer using LIP-1 sensors (Nicoya, SEN-AU-100-10-LIP1). The sensor surface was prepared for PM-mimetic vesicle immobilization with injections of 20 mM CHAPS (Nicoya, LIP-RK) at 150 μL/min. PM-mimetic vesicles (50 µg /mL) were flowed over the chip surface at 20 μL/min until immobilization was detected (600 AU minimum). An injection of 0.1% w/v bovine serum albumin (BSA, Fisher Bioreagents, BP9703100) in HBS was used to prevent non-specific protein binding to the sensor surface. Purified protein samples in HBS were flowed over the immobilized PM-mimetic vesicle at increasing concentrations. Injection of protein onto a CHAPS treated but PM-mimetic vesicle free sensor chip was used as a negative control. One-to-one kinetic models of interactions were analyzed in TraceDrawer 1.6.1 (LigandTracer).

### Cell culture

HEK293T cells were maintained in DMEM (Life Technologies, 4.5 g/L glucose, 110 mg/L pyruvate, 11960069), supplemented with 10% (v/v) Fetal Bovine Serum (FBS, Cytiva HyClone, SH3039603), and 1,000 U/mL penicillin/streptomycin (Gibco, 15140122). Inhibition of cholesterol synthesis was performed as previously published (Anderson et al., 2021). Briefly, after 48 h growth in 10% FBS containing media, cells were rinsed twice in serum-free DMEM and cultured under cholesterol depleted conditions in 7.5% lipoprotein deficient serum (LPDS) for 48 h with or without small molecule inhibitors AY9944 (2.5 µM, Cayman Chemical, 14611; *DHCR7* inhibitor). Acute sterol depletion was achieved by 1 h incubation with 5 mM MβCD (Alfa Aesar, J66847, 1303.31 g/mol) at 37°C. The 293T DVL2-mEGFP KI cell line was a kind gift from Dr. Xi He (Harvard University) (Ma et al., 2020).

### Human iPSC culture

Feeder-free human iPSCs were maintained in mTeSR1 (Stem Cell Technologies) on hESC-qualified Matrigel (BD Biosciences) coated culture dishes at 37°C in humidified 5% CO2 and 21% O2. Human iPSCs utilized for neural stem cell derivation were maintained on inactivated mouse embryonic fibroblasts (MEFs; Global Stem) in iPSC medium: DMEM/F12 base, 2 mM l-glutamine, 20% Knockout Serum Replacement (KOSR), 1% non-essential amino acids (NEAA), 1,000 units/ml Penicillin-streptomycin, 100 μM β-mercaptoethanol (Life Technologies), and basic fibroblast growth factor (bFGF; 7.5 ng/ml; Miltenyi Biotec). All iPSC cultures were passaged with Dispase (1 mg/ml; Stem Cell Technologies). Cultures were routinely analyzed by PCR for mycoplasma contamination/DNA (Universal Mycoplasma Detection Kit; ATCC). All cell culture reagents were batch tested for maintenance of hiPSC morphology and culture growth.

All research involving hiPSCs was approved by the Institutional Biosafety Committee at Sanford Research (approval no. 2015101). Clinical sample collection was approved by the Institutional Review Board of the Eunice Kennedy Shriver National Institute of Child Health and Human Development. Permission from guardians and consent, when possible, were obtained from participants. Clinical studies under which fibroblasts were collected for iPSC generation are posted on ClinicalTrials.gov (NCT00001721, NCT00046202 and NCT00344331).

### Neural stem cell differentiation of iPSCs

For generation of neural stem cell lines, embryoid bodies (EBs) were generated in iPSC media plus 20 ng/ml bFGF using AggreWell 800 dishes. After 24 h of culture, media was changed to neural induction media: DMEM (no glutamine, no sodium pyruvate) basal, N2A supplement, B27 with Vitamin A, epidermal growth factor (EGF; 20 ng/mL), bFGF (10 ng/mL), LDN193189 (100 nM), 2 mM L-glutamine, and 1,000 units/ml penicillin-streptomycin (Life Technologies).

After an additional 5 d, neuralized EBs were plated to poly-L-ornithine (20 µg/mL) and laminin (10 µg/ml)-coated dishes for neural rosette formation for 3-5 d. Rosettes were manually isolated, cultured until confluency, and passaged as neural stem cell (NSC) lines with Accutase (Life Technologies). NSCs were validated for expression of mouse anti-human Nestin (1:2,000; Millipore; MAB5326), rabbit-anti Sox2 (1:400; Cell Signaling; D6D9), and absence of mouse anti–βIII-tubulin (1:2,000; Millipore; MAB1637). Primary antibodies were detected with donkey anti-rabbit 555-IgG (1:500; Life Technologies; A31572) or goat anti-mouse 488-IgG (1:500; Life Technologies; A11001).

### Animals and housing

All animal work was reviewed and approved by the Institutional Animal Care and Use Committee at Sanford Research (protocol #186-09-24C). Mice were maintained on a C57Bl/6 background in normal light-dark cycle of 12 h/12 h with free access to standard chow and water. Littermates of both sexes housed together were used for all experiments. *Dhcr7*^Δ3-5/T93M^ mice were generated by mating *Dhcr7*^WT/Δ3-5^ with *Dhcr7*^T93M/T93M^ mice, both kindly provided by Dr. Forbes Porter, *Eunice Kennedy Shriver* National Institute of Child Health and Human Development. These heterozygous mice carry deletions of *Dhcr7* exons III, IV and part of V, and a dinucleotide mutation in codon 89 (Correa-Cerro et al., 2006; Wassif et al., 2001). As reported previously (Correa-Cerro et al., 2006; Freel et al., 2022), *Dhcr7*^WT/T93M^ mice were used as experimental controls for *Dhcr7*^Δ3-5/T93M^ mice.

### Preparation of lipoprotein-deficient serum

To evaluate the impact of cholesterol depletion on cultured cells, sterols, triglycerides, and other neutral lipids were depleted from fetal bovine serum (FBS) as described (Anderson et al., 2021; Francis et al., 2016). Ethylenediamine tetraacetate (EDTA, 0.1 mg, Invitrogen, 15-575-020) was added per 50 mL initial FBS. Phase separation of lipids from the serum was performed using an organic phase of diisopropyl ether: n-butanol (3:2; Sigma Aldrich, 296856, B7906, respectively). Organic and aqueous phases were allowed to mix at room temperature with continuous stirring in the dark for 1 h. The aqueous layer was collected by centrifugation, lyophilized, and resuspended in molecular grade water. Insulin-Transferrin-Selenium-G supplement (ITS-G, Gibco, 41-400-045) was added followed by filter-sterilization. Aliquots were stored at −20° C for further use. Each lot of lipoprotein deficient serum (LPDS) was validated for effect by GC-MS analysis of cholesterol levels in serum alone and in cells cultured with LPDS. LPDS was substituted for FBS at 7.5% in media to induce *de novo* cholesterol biosynthesis.

### Sterol loading of MβCD

Cells were treated with methyl-β-cyclodextrin (MβCD, Alfa Aesar, J66847) alone or complexed with cholesterol or other sterols as published (Anderson et al., 2021). For MβCD-sterol complex formation, 50 mg/mL sterol stocks (desmosterol, lathosterol, 7DHC, or cholesterol; Avanti Polar Lipids, 700060, 700069, 700066, 700000, respectively) were prepared in 1:1 chloroform: methanol. Following previously determined optimal loading conditions (Anderson et al.,2021), a stoichiometry of 1:7 molar ratio was followed to conjugate MβCD-sterol complexes. The desired amount of sterol was dried under nitrogen flow and resuspended in 5 mM MβCD (Alfa Aesar, 1303.31 g/mol) in serum-free DMEM by 5-minute probe sonication followed by agitation at 37°C for 8 h. Prior to each experiment, the solution is filtered through a 0.22 µm membrane for sterilization and removal of crystals. Rescue experiments consist of incubation in the complex solution for 1 h at 37°C supplemented with 0.5% BSA, followed by removal, wash, and addition of 7.5% LPDS to limit MβCD toxicity.

### Quantitative analysis of sterols by GC-MS

Quantitative sterol analysis of cell pellets or PM-mimetic vesicles was performed as described (Anderson et al., 2021; Francis et al., 2016). TMS derivatives of sterols exhibiting <3% abundance were excluded from analysis. For sterol quantitation, abundance was normalized to both the internal standard (coprostanol, Cayman Chemical, 26764) and protein concentration measured by BCA assay. For PM-mimetic vesicles, sterol abundance was normalized to the internal standard only. Analysis was performed using MassHunter software. Identification of TMS ethers of natural sterols was determined through comparison to commercially available standards, as well as comparison to MS spectra through the National Institute of Standards and Technologies Standard Reference Database when annotated. Retention times and mass to charge (m/z) ratios were described previously (Anderson et al., 2021). Statistical significance was calculated in GraphPad Prism software.

### Subcellular fractionation

To obtain subcellular fractions by differential centrifugation, HEK293T cells were plated out in 60 mm dishes coated with 0.1% gelatin (Millipore, SF008). After treatment and/or rescue, cells were gently scraped into 250 μL ice-cold fractionation buffer [20 mM HEPES (pH 7.4, Fisher Scientific, BP310-100), 10 mM KCl (Milipore, 529552), 2 mM MgCl_2_ (Sigma-Aldrich, 208337), 1 mM EDTA, 1 mM EGTA (Sigma-Aldrich, E8145), 1 mM DTT (Sigma-Aldrich, D0632), protease inhibitor cocktail (Fisher Scientific, PIA32955)], and incubated on ice for 15 min. All procedures were performed at 4° C. A sterile 1 mL syringe was used to gently pass the cell suspension through a 27-gauge needle 10 times followed by 20 min incubation on ice. Samples were centrifuged at 720 x g (3,000 rpm) for 5 min. The supernatant (S1) was saved for membrane protein isolation. The nuclear pellet (P1) was washed twice with 500 μL fractionation buffer and dispersed with a pipette to resuspend in 100 μL fractionation buffer with 0.1% Triton-X-100. The suspension was then passed through a 27-gauge needle 10 times followed by centrifugation at 24,000g at 4° C for 10 minutes. The resultant nuclear protein in supernatant was stored at −20° C. To obtain membrane fraction, the initial supernatant S1 was centrifuged at 8,000 rpm (10,000 x g) for 5 min. Resultant supernatant S2 was further centrifuged in a WX-80 Ultra centrifuge (ThermoFisher Scientific) at 100,000 x g for 1 h. Supernatant S3 is the cytosolic fraction. Pellet P2 was washed twice in fractionation buffer and resuspended in 400 μL of fractionation buffer by passing through a 25-gauge needle. Centrifugation for 45 min at 100,000 x g for 1 h yielded a membrane pellet P3. The membrane pellet was washed twice in fractionation buffer and resuspended in 50-100 μL of fractionation buffer supplemented with 0.1% Triton-X-100. Protein concentration was measured with standard BCA assay according to manufacturer’s instructions (Fisher Sci, PI23227).

### Co-immunoprecipitation

For subcellular co-immunoprecipitation (Co-IP), organelle fractions were resuspended in a non-denaturing lysis buffer (20 mM Tris HCl pH 8, 137 mM NaCl, 1% Nonidet P-40 (NP-40) and 2 mM EDTA, supplemented by protease inhibitors). The starting protein was maintained at 500 μg in 0.2-0.5 μL. The protein fraction was precleared for 1 h with 50 μL of normal rabbit serum (Jackson ImmunoResearch, 011-000-120) and 50 μL of agarose A/G bead slurry (Pierce™ Protein A/G Agarose, Fisher Scientific, PI20421) at 4°C with gentle agitation. The reaction was centrifuged at 14,000 x g at 4°C for 10 min. The bead pellet was discarded and supernatant was used for immunoprecipitation. The protein sample was incubated with appropriate primary antibody (rabbit anti-DVL2 polyclonal antibody [N1N3], GeneTex, GTX111156) for 12 h at 4°C, with gentle agitation. 50 μL of blocked and washed (0.1% BSA in PBS, 1 h) beads were added to the reaction and incubated at 4°C under rotary agitation for 4 h. The supernatant was removed from the beads by centrifugation and after three rinses in wash buffer (10 mM Tris pH 7.4, 1 mM EDTA, 1 mM EGTA; pH 8.0, 15 0mM NaCl,1% Triton X-100, 0.2 mM sodium orthovanadate [Sigma-Aldrich, S6508], protease inhibitor cocktail), protein was eluted from the beads in 50 μL 2X dye-less Laemmli buffer (65.8 mM Tris-HCl, pH 6.8, 2.1% SDS, 26.3% (w/v) glycerol, without DTT to minimize antibody co-elution) at 50° C for 10 min. Samples were run in SDS-PAGE and western blotting was performed to detect the binding of candidate proteins to DVL2.

### Immunocytochemistry

For immunocytochemistry (ICC), cells were plated on 12 mm glass coverslips and subjected to appropriate treatments prior to staining. Cells were fixed in 4% paraformaldehyde for 20 min, rinsed three times in PBS, and permeabilized in 0.2% Triton X-100 for 20 min, followed by blocking in 5% goat or donkey serum (Jackson ImmunoResearch, 017-000-121) in 0.1% Triton X-100 at RT for 1 h. Primary antibody incubation in appropriate dilution was performed overnight in blocking buffer at 4°C. Primary antibody (rabbit anti-DVL2, N1N3, GeneTex, GTX111156; 1:1,000) was visualized with Alexa Fluor™ conjugated secondary antibodies (1:500, Life Technologies). Hoechst 33342 nuclear counterstain (Invitrogen, H3570, 1:10,000) or Alexa Fluor™ 647 Phalloidin (Invitrogen, A22287, 1:400) were incubated with the secondary antibodies where indicated. Images were captured using a Nikon A1R resonant scanning multispectral confocal microscope (Nikon Instruments, Inc. Melville, NY) Nikon Ti Perfect Focus system, and NIS-Elements analysis software (Nikon).

### TurboID proximity-dependent protein-protein interaction screening

Human DVL2 was cloned into pBabe-puro-TurboID following previously published methods (May et al., 2020). HEK293T cells were used to generate stable HsDVL2-turboID expressing lines (May et al., 2020). TurboID expression was confirmed using immunofluorescence and western blots (**Figure S6A-B**). Cells were propagated in standard FBS-containing media. To quantify the impact of sterol availability on DVL2 binding partners, cells were cultured in media containing 7.5% LPDS and AY9944 for 48 h. Cells were treated with 50 µM biotin for 18 h prior to isolation. Turbo-ID pulldowns (**Figure S6A-B**), LC-MS fractionation, peptide identification and analyses (**Table S5**) were performed as previously described (May et al., 2020; Sears et al., 2019). Protein-protein interaction networks were generated in STRING database and analyzed for enrichment in Cytoscape software. Network clustering has been performed in Cytoscape. Reactome database search, GO enrichment and KEGG pathway analysis are performed for data analysis and visualization.

### Statistical analysis

All statistical analyses were performed using GraphPad Prism 8.0.2 (GraphPad Software, Inc., CA, US). Equal variance was assumed. Data were analyzed by one-way ANOVA, two-way ANOVA, or student’s t-test with post hoc tests for multiple comparisons relative to controls. p < 0.05 was accepted as significant. Specific statistical tests and details for each experiment can be found within the corresponding figure legends.

## Data availability

Datasets generated in this study are available from the corresponding author upon request.

