## Supplemental Information_Sengupta et al for "Dishevelled localization and function are differentially regulated by structurally distinct sterols"

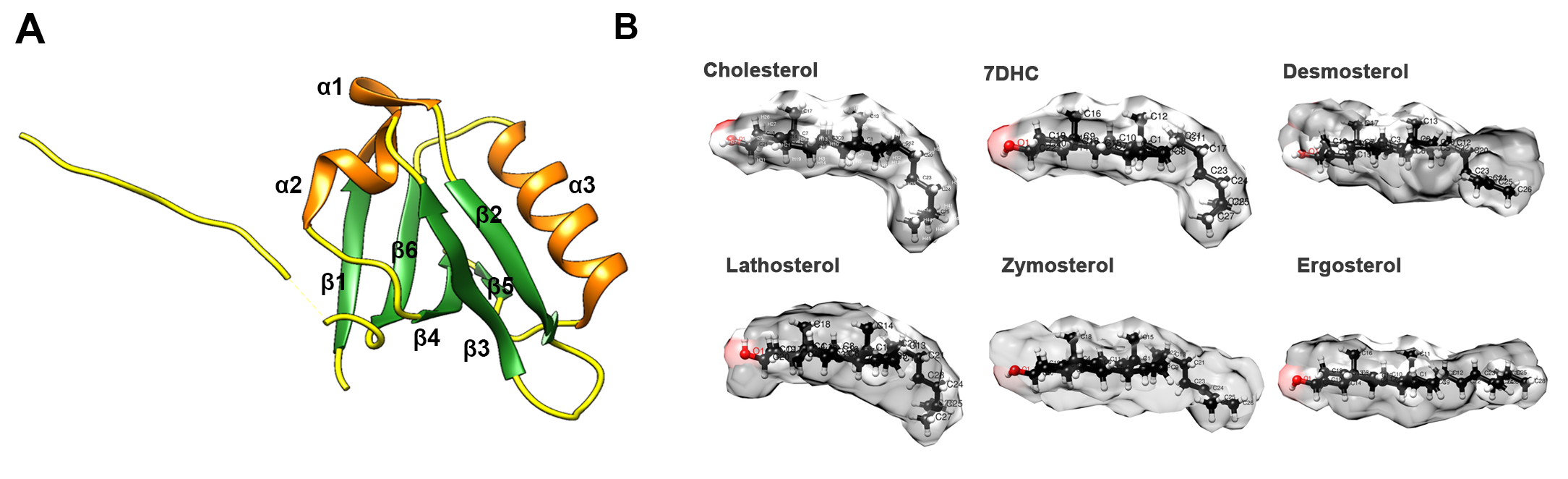

**Figure S1. Structural comparison of cholesterol and alternate sterols of biological interest. Related to Figure 1.**

(A) Three-dimensional ribbon structure of the PDZ domain of human DVL2 (PDB ID: 3CBZ), colored and numbered according to the secondary structure. Coils = yellow; β-strands = green; α-helices = orange.

(B) Three-dimensional, space filling conformation of sterols used in this study. The C3 polar headgroup is colored red.

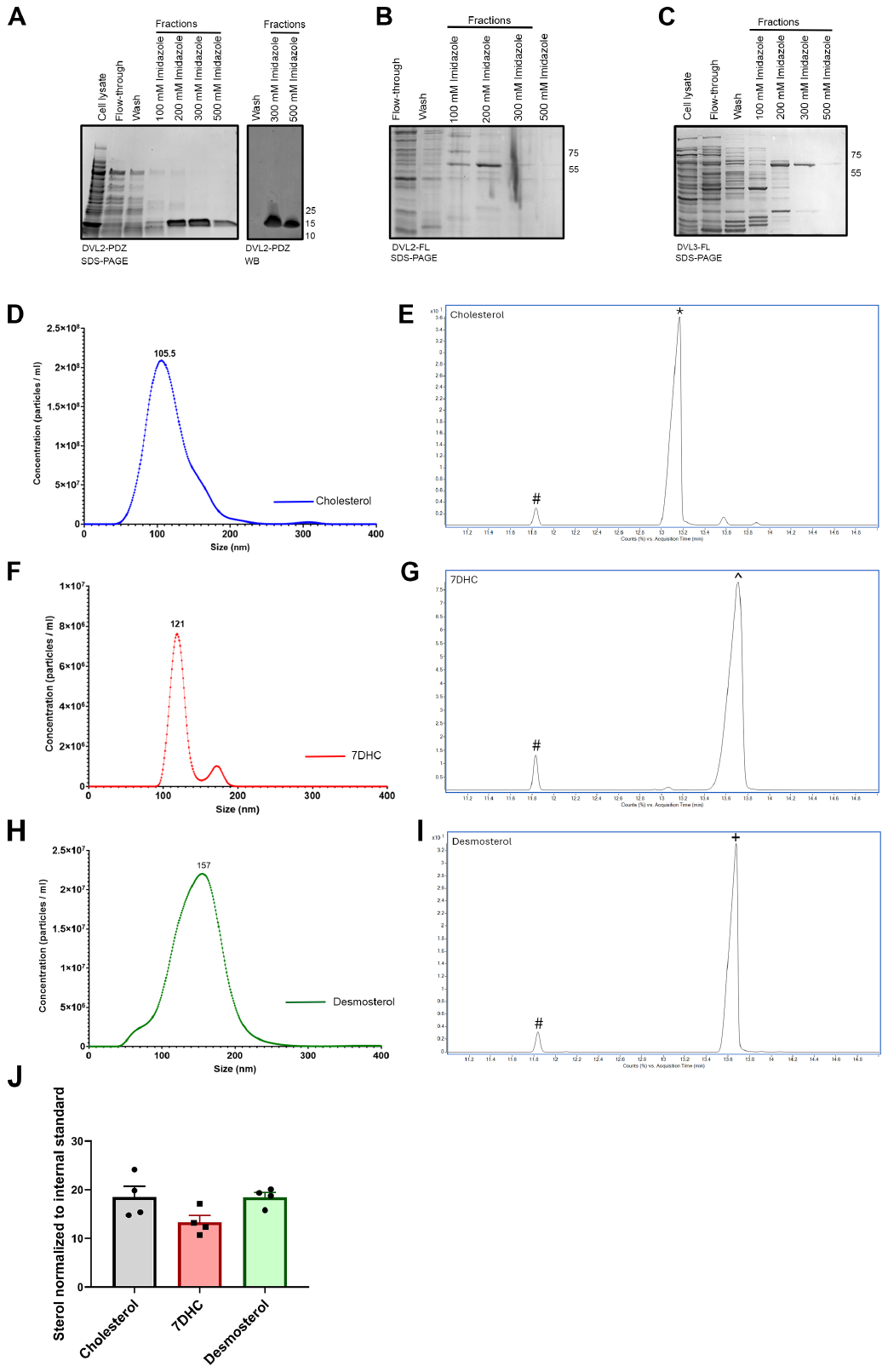

**Figure S2. Validation of DVL protein purification and PM-mimetic vesicle preparation for protein-lipid interaction analyses. Related to Figure 2.**

(A) Ni-NTA resin purification of His-tagged DVL2-PDZ protein (~15 kDa); left = SDS-PAGE of cell lysate and fractions from purification; right = western blot of concentrated fractions using α-His antibody (1:5000).

(B) SDS-PAGE of Ni-NTA resin purified fractions of His-tagged DVL2-FL (~58 kDa), showing cell lysate, column flow-through, wash, and fractions eluted at increasing concentrations of imidazole (100-500 mM). Majority of protein was eluted at 200 mM imidazole.

(C) SDS-PAGE of Ni-NTA resin purified fractions of His-tagged DVL3-FL (~57 kDa), showing cell lysate, column flow-through, wash and fractions eluted at increasing concentrations of imidazole (100-500 mM). Majority of protein was eluted at 200-300 mM imidazole.

(D, F, H) Representative particle size analysis of cholesterol-incorporated PM-mimetic vesicles (cholesterol, 7DHC, and desmosterol) detected by Nanosight.

(E, G, I) Representative GC-MS chromatograms for trimethylsilyl derivatized sterols following incorporation into PM-mimetic vesicles (cholesterol, 7DHC, and desmosterol).

(J) Quantitation of cholesterol and representative alternate sterol incorporation into PM-mimetic vesicles measured by GC-MS. N = 4 independent preparations ± SEM. # = coprostanol (internal sterol standard), * = cholesterol, ∧ = 7-dehydrocholesterol, +=desmosterol.

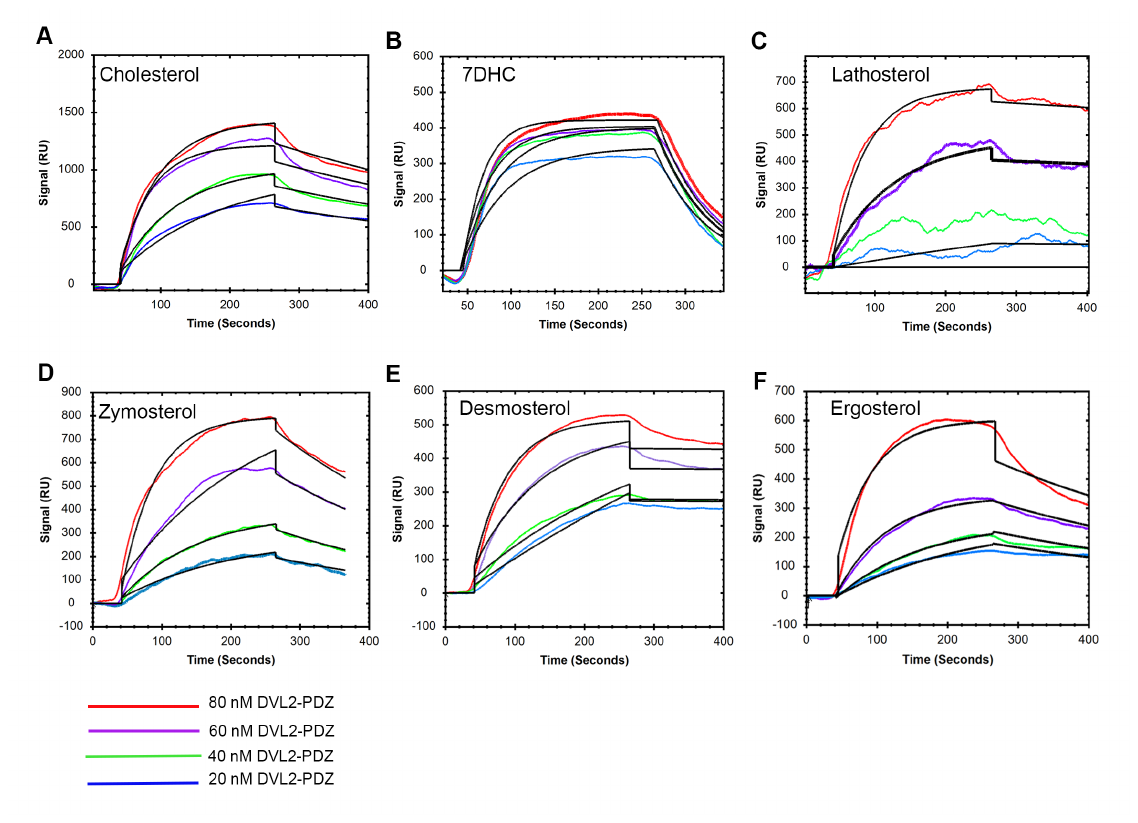
 **Figure S3. Representative SPR sensorgrams fitted to 1:1 binding model for respective sterol-enriched PM-mimetic vesicles. Related to Figure 3**.

(A-F) Representative sensorgrams with 1:1 binding curve fittings of DVL2-PDZ binding to PM-mimetic vesicles containing cholesterol (A), 7DHC (B), lathosterol (C), zymosterol (D), desmosterol (E), or ergosterol (F). Fitted sensorgrams were generated using TraceDrawer software.

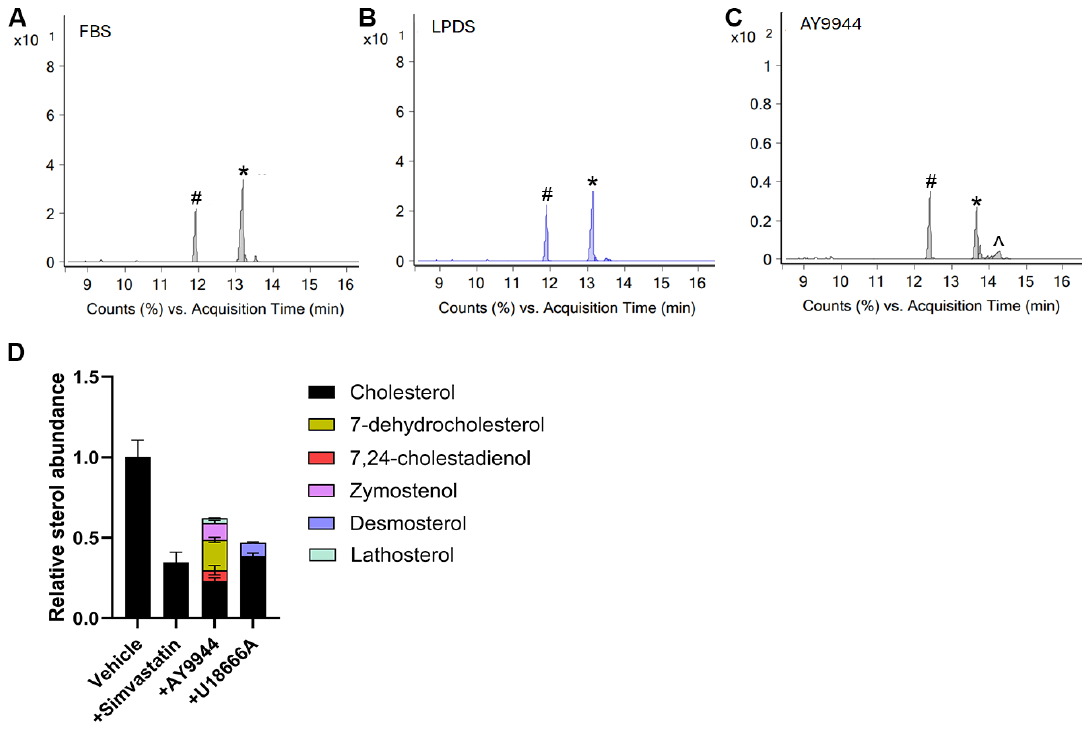
**Figure S4. GC-MS analysis of HEK293T cells demonstrate characteristic sterol accumulation pattern following cholesterol depletion or AY9944 treatment. Related to Figure 4.**

(A-C) Representative chromatograms of HEK293T cells cultured in FBS or LPDS media and/or treated with AY9944.

(D) GC/MS analyses of cells in cholesterol depleted LPDS (vehicle), Simvastatin, AY9944, or U18666A treated conditions. Data represents the mean±SEM, N=3 biological replicates. # = coprostanol (internal standard), * = cholesterol, ∧ = 7- dehydrocholesterol.

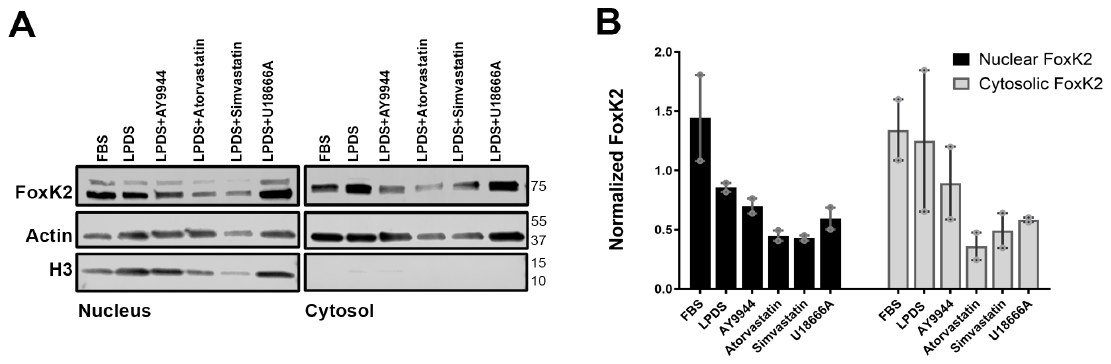

**Figure S5. Subcellular FoxK2 expression is not changed following cholesterol-targeted treatments. Related to Figure 5.**

(A) Western blot analyses of nuclear and cytosolic Foxk2 expression under various sterol-targeted conditions.

(B) Quantification of nuclear and cytosolic fractions demonstrate that FoxK2 expression does not significantly change following sterol targeted treatments. N = 2 independent experiments, two-way ANOVA with Dunnett’s post-hoc multiple comparison test, *p < 0.05, **p < 0.01, ***p < 0.001, ***p < 0.0001 for all statistical tests.

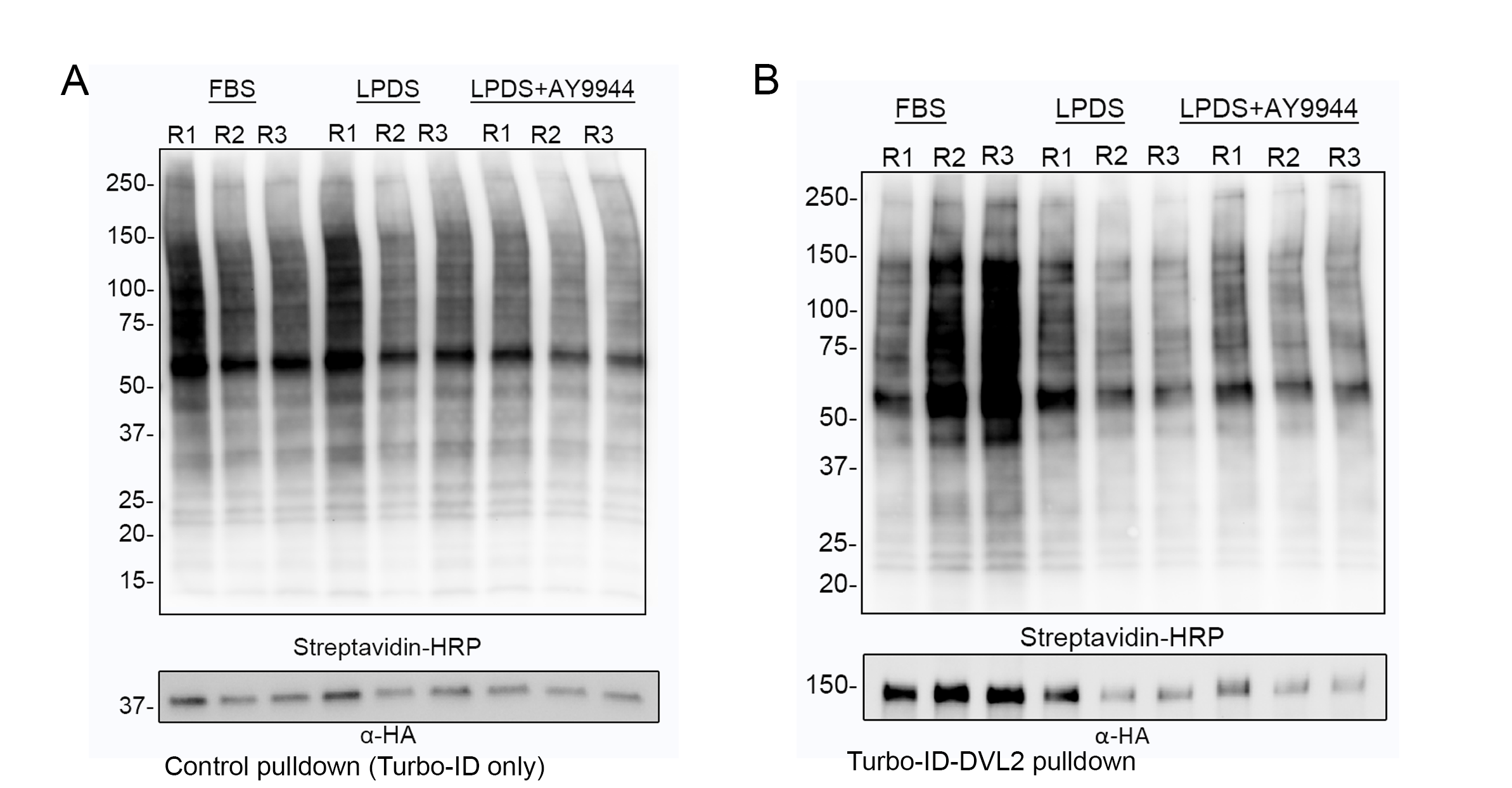

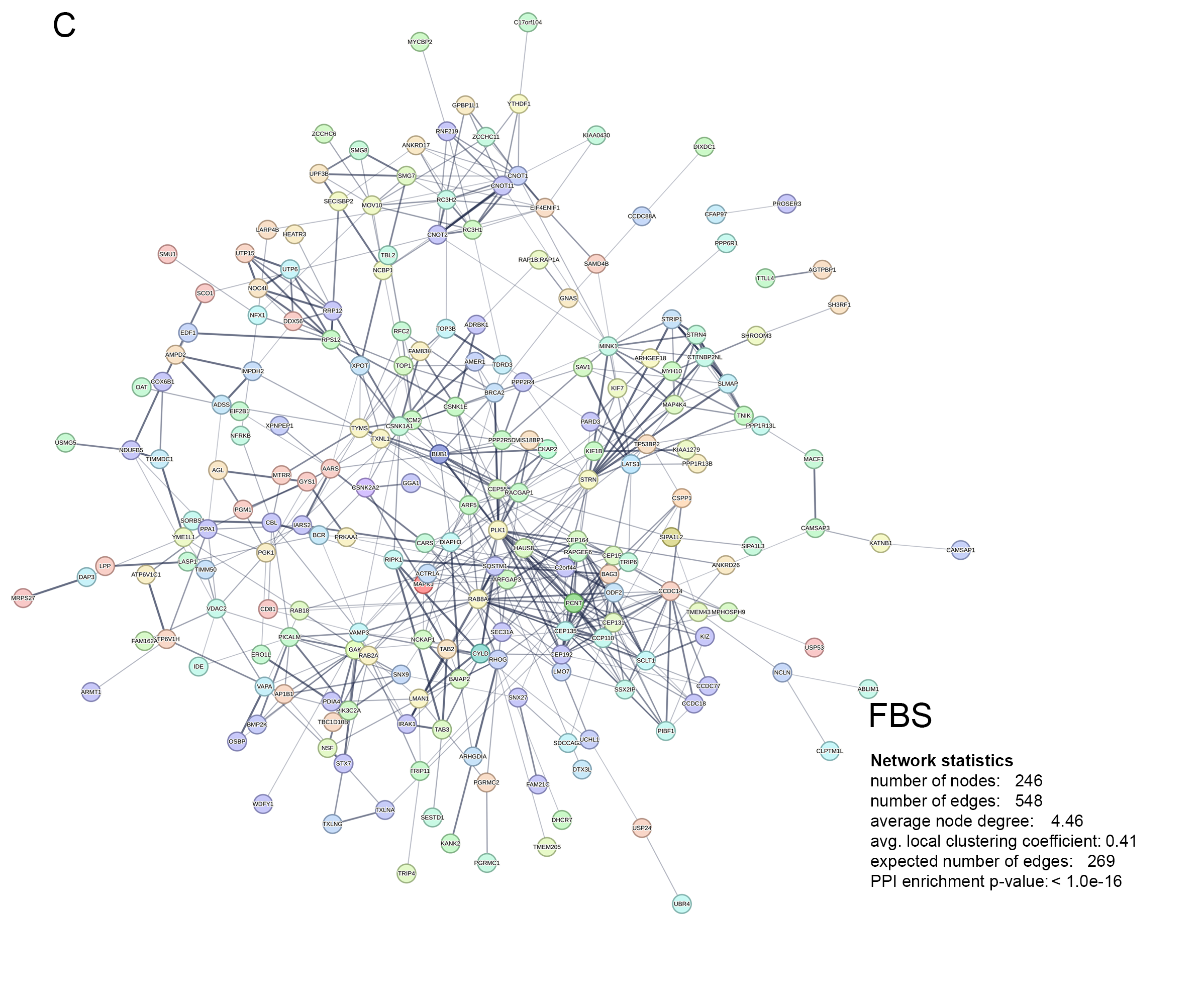

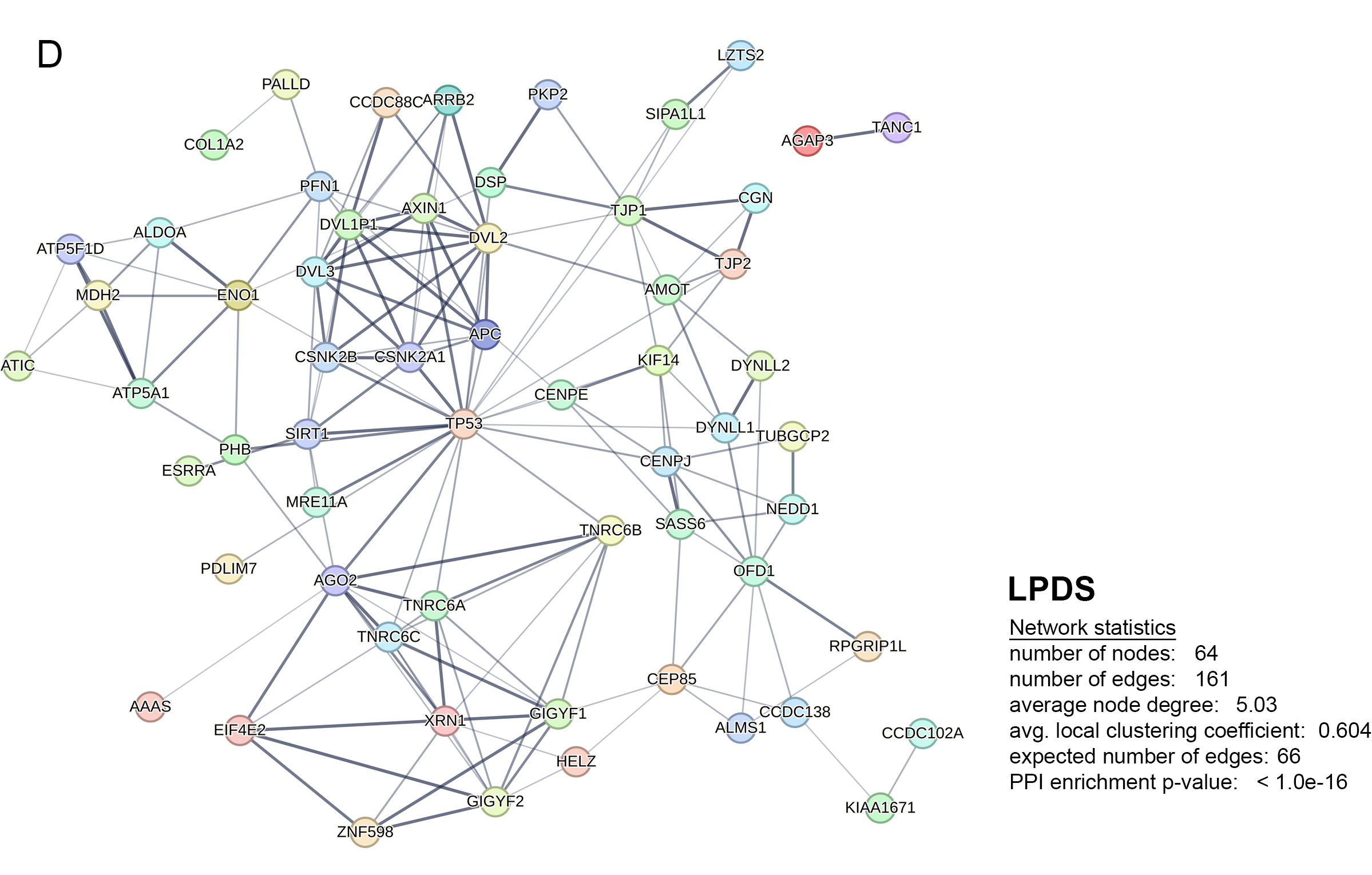

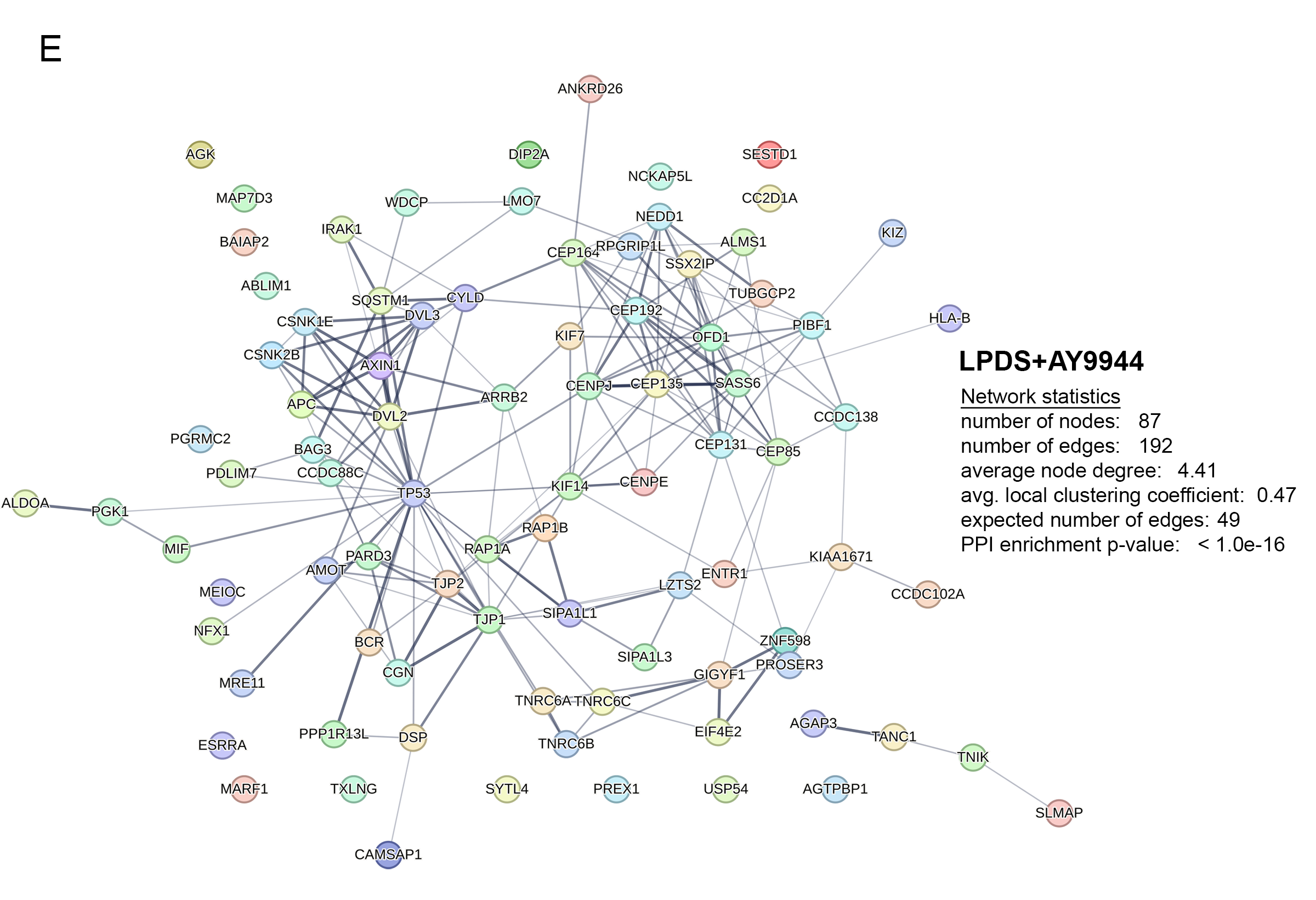
**Figure S6. Pulldown validation and protein-protein interaction analyses resulting from TurboID-DVL2 assays. Related to Figure 6.**

(A, B) Post-pulldown immunoblot (WB) detection of enriched biotinylated proteins with streptavidin-HRP in the Turbo-ID-DVL2 proximity-based ligation assay. R1-R3 are the biological replicates under each treatment. The DVL2-Turbo-ID assays show DVL2-TurboID-HA fusion protein detected by anti-HA antibody (B, lower panel).

(C-E). STRING analysis identification of enrichment networks from DVL2 protein-protein interactions detected within each treatment. LPDS+AY9944 (87 nodes + 192 edges) compared to LPDS (64 nodes + 161 edges) indicating complexity of network. Standard FBS culture (cells receiving cholesterol rich media) exhibited higher complexity (246 nodes + 548 edges).

| Sterol | LogP (negative measure of hydrophobicity) | Topological polar surface area(A°) | Van der Waals molecular volume | Hydrogen bond donor/  acceptor | Biological and clinical significance |
| --- | --- | --- | --- | --- | --- |
| Cholesterol | 7.68 | 20.23 | 432.37 | 1/1 | Broad mammalian expression; critical to membrane function, lipid ordering; high levels in cardiovascular diseases, inflammation, cancers (1-4) |
| 7-dehydrocholesterol | 7.60 | 20.23 | 429.73 | 1/1 | Expressed in keratinocytes; accumulates in Smith-Lemli-Opitz syndrome (5-7) |
| Desmosterol | 7.60 | 20.23 | 429.73 | 1/1 | Expressed in astrocytes, sperm, macrophages, Desmosterolosis, inflammation (8-13)  ­ |
| Lathosterol | 7.68 | 20.23 | 432.37 | 1/1 | Expressed in brain, increased in Lathosterolosis, adenomas; decreased in aging brain, colorectal cancer (14-17) |
| Ergosterol | 7.62 | 20.23 | 444.39 | 1/1 | Predominant sterol within fungi (18, 19) |
| Zymosterol | 7.74 | 20.23 | 429.73 | 1/1 | Increased in Huntington’s disease, Conradi-Hunermann syndrome (20, 21) |

**Table S1. Physical properties and biological significance of sterols used in this study.** Structurally disparate sterols were chosen based on differences in ring and hydrocarbon tail saturation, as well as relevance to mammalian biology and human disease. Hydrophobicity and polarity measures were obtained from PubChem (https://pubchem.ncbi.nlm.nih.gov). Van der Waals and hydrogen bonding values for individual sterols were obtained from Lipid Maps (https://lipidmaps.org). **Related to Figure 1.**

| **Target sterol** | **Predicted binding energy (kcal/mol)** | **RMSD/lower bound** | **RMSD/upper bound** |
| --- | --- | --- | --- |
| Cholesterol | -7.3 | 0 | 0 |
| 7DHC | -6.4 | 0 | 0 |
| Desmosterol | -5.7 | 0 | 0 |
| Lathosterol | -5.3 | 0 | 0 |
| Zymosterol | -6.8 | 0 | 0 |
| Ergosterol | -3.2 | 0 | 0 |
| 7DHC in “cholesterol orientation” | -6.1 | 2.082 | 9.136 |
| Ergosterol in “cholesterol orientation” | -6.4 | 16.299 | 18.545 |

**Table S2. Predicted binding energies of selected sterols to DVL2-PDZ.** Binding of individual sterols to DVL2-PDZ domain protein (PDV ID: 3CBZ) were calculated by AutoDock Vina software (https://vina.scripps.edu) using default settings. Results shown for root mean square deviation (RMSD) values <2.0 represent predicted binding in the ‘optimal’ sterol orientation identified by AutoDock Vina. To mimic cholesterol’s orientation regarding DVL2-PDZ binding, predicted energies and RMSD values for 7DHC and ergosterol >2.0 are also presented. **Related to Figure 1.**

| **Human iPSC line** | **Source subject, tissue** | **Clinical phenotype** | ***DHCR7* mutation** |
| --- | --- | --- | --- |
| BJ 3F1 | BJ neonatal fibroblasts (ATCC) | Unaffected control | None |
| CWI 4F2 | SLOS-029 patient fibroblasts (NIH) | Smith-Lemli-Opitz syndrome | p.T93M/c.964-1G>C |
| dCas9-KRAB WTC-11 | Dermal fibroblasts | Constitutive dCas9 expression from CLYBL locus | None |

**Table S3**. **Human iPSC lines utilized in this study.** Cell lines were previously described (22-24)**. Related to Figure 7.**

| **Antibody used** | **Supplier** | **Catalog number** | **Concentration used** |
| --- | --- | --- | --- |
| His-Tag, rabbit pAb | Cell Signaling Technology | #2365 | 1:1000 |
| β-Actin (8H10D10), mouse mAb | Cell Signaling Technology | #3700 | 1:1000 |
| Anti-Actin antibody, mouse mAb | Sigma Aldrich | #A3853 | 1:1000 |
| β-Catenin (D10A8), XP® rabbit mAb | Cell Signaling Technology | #8480 | 1:1000 |
| Dishevelled 2 antibody [N1N3], rabbit pAb | GeneTex | # GTX111156 | 1:500 |
| FOXK2 antibody [C3], C-term rabbit pAb | GeneTex | #GTX104848 | 1:1000 |
| Histone H3, mouse mAb | LI-COR | #926-42218 | 1:1000 |
| V-ATPase B1/2 Antibody (F-6), mouse mAb | Santa Cruz | #sc-55544 | 1:200 |
| p53 (1C12), mouse mAb | Cell Signaling Technology | #2524T | 1:1000 |
| Lamin A antibody(C-3), mouse mAb | Santa Cruz | #sc-518013 | 1:1000 |
| GAPDH antibody, mouse mAb | ThermoFisher | #AM4300 | 1:1000 |

**Table S4. Antibodies and concentrations utilized in this study. Related to Figures 2, 4, 5, 6, and 7.**

**Table S5. DVL2 interaction proteins identified by mass spectrometry analyses of proximity-based labeling in HEK293T cells. Related to Figure 6.**

(A) Mass spectrometry data for identification of DVL2 proximity interactions using TurboID-DVL2 in FBS, LPDS, and LPDS+AY9944 conditions. TurboID-only was used separately for each condition.

(B) Analysis of TurboID-DVL2 in FBS conditions.

(C) Analysis of TurboID-DVL2 in LPDS conditions.

(D) Analysis of TurboID-DVL2 in LPDS+AY9944 conditions.

(E) TurboID identified candidate interactors of DVL2 in HEK293T cells by condition
